# Tinkering loci: hubs of gene birth in an animal genome

**DOI:** 10.64898/2026.08.09.743762

**Authors:** Caroline M. Weisman

## Abstract

How does evolution create new things? A key strategy is “tinkering”: changing and re-using existing genetic components in new ways. Tinkering is generally conceived of as unpatterned, occurring without spatial or other kinds of structure within the genome. Here, I report “tinkering loci”: distinct genomic regions that significantly accelerate gene birth by tinkering. This occurs because tinkering loci accumulate unusually high concentrations of duplicated gene fragments from around the genome, which they then re-transcribe. Because they bring previously unrelated genes into proximity much more often than expected by chance, they are especially strong accelerants of composite gene creation. I find that tinkering loci are common in *Drosophila* genomes; vary in number and activity over a few million years; are driven by a standard mutational mechanism acting at all major kinds of transposon sequences; and disperse, as well as receive, duplicates, including their functional products. Tinkering loci demonstrate the possibility of nonrandom genomic patterning of mutations that meaningfully shapes the rates and consequences of molecular innovation. Because these patterns appear at least in part to be themselves encoded in the genome via transposons, which are often genomically abundant and highly dynamic, they suggest a rich and evolvable dimension of genomes that could influence the tempo and mode of animal evolution.

## Introduction

Genes are fundamental pieces from which organisms are built. Understanding the mechanisms that generate new ones is crucial for a robust understanding of evolution. It may also provide insight into mechanisms underlying other types of genetic innovation.

François Jacob famously described evolution as a “tinkerer” who innovates by altering genetic components that it has already made and re-using them in new ways^1^. This powerful strategy accelerates evolution by taking advantage of eons of past work and puts within reach inventions whose creation “from scratch” would be prohibitively slow. There is now substantial evidence that tinkering is central to evolutionary innovation, especially in complex multicellular taxa^2–5^. This is particularly well-documented for the creation of new genes, which can be formed by the complete or partial duplication of existing genes^6^ and by “shuffling,” an umbrella term in which pieces from different existing genes are combined into new ones, including chimerism^7^, exon shuffling^8^, and domain shuffling^9^. These tinkering processes are generally described in terms of individual mutations distributed more or less randomly throughout the genome: the effects of unpatterned mutations, with little influence of the overall architecture of the genomes in which they occur^10–12^.

Despite the evolutionary importance of tinkering, we still lack a complete picture of the mechanisms by which it is carried out. For example, “orphan” genes, whose origins cannot be identified, are pervasive^13–19^, suggesting a missing source of gene birth. (These are sometimes interpreted as having originated “de novo” from noncoding sequence, but this is often an unreliable inference made based on an absence of evidence^19–23.)^ Beyond new genes in particular, many have proposed that known mutational mechanisms are not, by themselves, enough to explain how tinkering produces the diverse and complex innovations that we observe. A broad suggestion often converged upon in different forms is that there are unappreciated patterns or structure in some aspect of how these mutations are deployed that enable them to more readily hit upon useful innovations^24–31^.

## Results

### Origin and spread of a young essential *Drosophila* gene at unusual genomic regions

As a case study approach to identifying mechanisms of gene birth, I investigated the origin of *goddard*, a well-studied and evolutionarily new gene in *Drosophila melanogaster*. It is essential for male fertility, but completely lacks detectable homologs outside of the family *Drosophilidae.* On this basis, it has been hypothesized to have originated “de novo” from noncoding DNA^32–34^.

For clues to its origin, I used available *Drosophilidae* genome sequences (Table S1) to examine the genomic region surrounding it, where I noticed unusual features (Methods-1). First, in *melanogaster* and most *Drosophila*, the *goddard* locus contains three apparently family-specific genes. Given locus size and the number of such genes in *melanogaster,* this is an unusually high density (p=6*10^-3^, Methods-2). Second, across the phylogeny, genes native to the locus are often rearranged and/or reoriented within it. Finally, the locus also contains many different complete and partial duplicate copies of existing genes. These duplicates are “<u>dispersed</u>”: the ancestral, conserved copies from which they are derived are not nearby, but in diverse and distant regions of the genome. In one species, *Chymomyza caudatula,* these include a large tandem array of a composite sequence, made from fragments of two unrelated genes that have here been placed immediately adjacent. One fragment derives from a homolog of *artemis*, another *melanogaster* gene that shares *goddard*’s unusual status as a young gene that has quickly become essential^35^. Examples of the locus in this and a few other species, not exhaustive of its diversity across *Drosophila*, are shown in Fig. 1A.

**Fig. 1:**
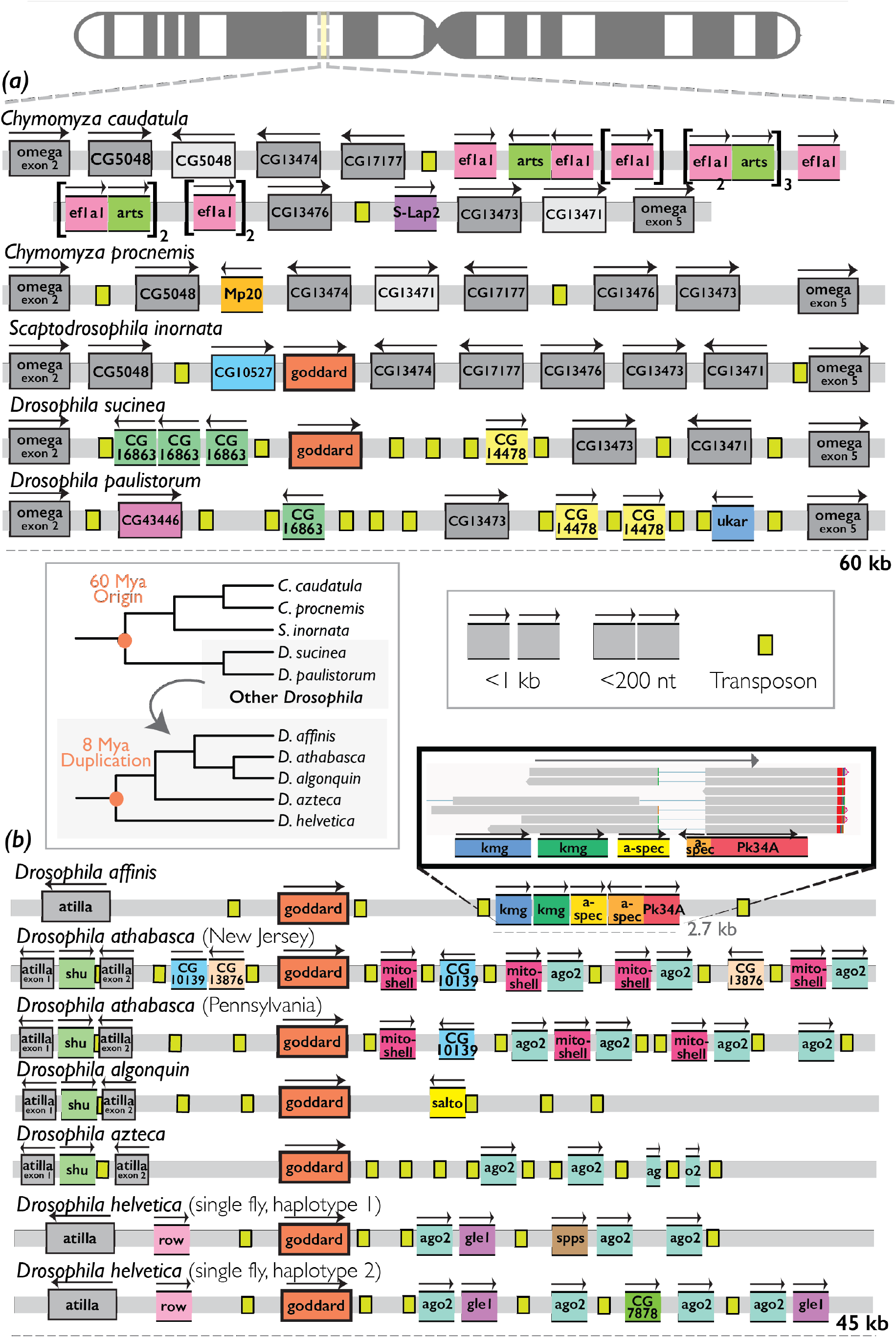
Origin and spread of *goddard* at unusual genomic regions. Figure depicts regions of the genome, as schematized by the chromosome cartoon at the top. Both panels: gray bars represent unannotated chromosomal sequence. Rectangles with intact black borders represent full-length genes (defined relative to the *melanogaster* sequence). Squares lacking black side borders represent gene fragments. Genes are labeled with the FlyBase^37^ abbreviation of the best *melanogaster* homolog gene name. Gray genes are conserved at the locus in *Drosophilidae*; lighter gray are those whose position or orientation is altered relative to other species. The red gene with a thicker black border is *goddard*. Other colored genes and fragments are specific to one or a few species; each unique color represents a unique fragment (e.g. all *ago2* fragments, in cyan, cover approximately the same region). Arrows above genes indicate strand. Small yellow-green boxes are transposons. For legibility, distances between elements are <u>not to scale,</u> but close spacing categories are represented according to the legend in the right-center of the figure. In the left-center of the figure are two phylogenies depicting the relationship between the species in the top subfigure, in the bottom subfigure, and the relationship of the two groups to one another. (**A**): The orthologous *goddard* locus in five *Drosophilidae. Chymomyza caudatula* is shown on two lines for ease of viewing because the number of features is especially large. *In paulistorum*, *goddard* has moved elsewhere in the genome; in *Chymomyza* species, it was likely lost after its origin. (**B**): Locus around a *goddard* duplicate specific to the *affinis* group. Inset: *affinis* Iso-Seq data showing fragments transcribed together on a “composite” transcript. Each gray bar is a single read; thin regions joining bars are spliced regions; red regions at the ends of transcripts are polyA tails.

I then noticed that in the “*affinis* group” of *Drosophila,* five species diverged by only ∼8 million years^36^, *goddard* has duplicated to a second locus with the same unusual properties (Fig. 1B).

This locus also contains many dispersed partial gene duplicates (hereafter “fragments”). There is substantial variation between species in which fragments are present, and fragments present in multiple species vary in their number and position. This variation persists even at shorter evolutionary distances: within a species (different *Drosophila athabasca* populations^38^) and an individual animal (two chromosomes from a single *Drosophila helvetica*^39^).

I performed long-read RNA sequencing with PacBio Iso-Seq on live *Drosophila affinis* (Methods-3), revealing that a dense complex of five ancestrally noncontiguous pieces from three genes is transcribed together, forming a novel “composite” transcript (Fig. 1B, inset).

### “Tinkering loci”: hubs of gene birth by tinkering

Tinkering is generally described in terms of individual mutations, more or less randomly distributed in the genome. In contrast, Figure 1 shows these mutations accumulating preferentially within kilobase-scale loci.

What effects might this unusual “mutational architecture” have? These loci accumulate gene duplicates dispersed from distant locations. Dispersed duplications are usually rare compared to “tandem” duplications, in which duplicates remain adjacent to their parent gene^40^. This usual bias works against gene birth: tandem duplicates less readily become functionally important than dispersed ones^41^, the latter possibly better able to contribute usefully in their novel regulatory environments^42^. Insofar as these loci significantly increase the dispersed duplication rate, they may accelerate gene birth.

These loci are also “hubs” that centralize duplicates from many different locations into a small region, resulting in a very high local concentration of originally distant genetic components. In classical shuffling, genes must attain proximity sufficient to create a new composite gene purely by chance, which is unlikely in large animal genomes. This seriously limits the rate at which successful products can be formed^43^. When it does happen, successful shuffling overwhelmingly occurs between genes that are already neighbors^44,45^, which excludes a huge fraction of the total combinations theoretically available to evolution. Hubs can overcome this barrier, enabling many combinations of genes that may otherwise have remained inaccessible.

Because of their potential to enhance gene birth by tinkering, I refer to these regions, and potential others like them, as “tinkering loci.”

### Tinkering loci are a general feature of *Drosophila* genomes

I first asked: are there other tinkering loci in *Drosophila*? I used their hallmark characteristic, an unusually high density of dispersed gene duplicates and fragments, to search for them systematically. Because gene fragments are excluded by most annotation methods^46,47^, I developed a pipeline to annotate gene fragments and duplicates (for brevity, hereafter jointly referred to as “fragments”) along with conserved genes (Methods-4). I applied this to five phylogenetically diverse *Drosophila* species: *virilis, grimshawi, repletoides, pseudoobscura, and affinis* (Fig. 2C). I then identified dispersed fragments in each genome (Methods-5). Their number varied >3x across species, highest (4180) in *affinis* and lowest (1189) in *grimshawi* (Fig. S1). If dispersed randomly through the genome, distances *x* between adjacent fragments would follow an exponential distribution, *e*^−*λ*x^, where *λ* is the inverse of their mean spacing. However, in their best fit to this null model, all species show an excess of fragments spaced closer than ∼10 kb (Fig. 2A), around the scale of the fragments in the exemplar tinkering loci (Fig. 1). In a simple alternative model, the genome has discrete tinkering loci in which fragments are more closely spaced than in the rest of the background genome (Fig. 2B). The overall distance distribution here becomes the mixture *π*_1_*e*^−*λ*1x^ + *π*_2_*e*^−*λ*2x^, where *λ*_1_,*λ*_2_ are the inverses of mean spacing in tinkering loci and background, and *π*_1_, *π*_2_ (*π*_1_+ *π*_2_ =1) the fractions of fragments in the two region types. This model fits the data significantly better in all species, while the single exponential better fits a negative control of randomly positioned fragments (Methods-6, Table S2), indicating quantitatively that fragments are spaced nonrandomly in the genome.

**Fig. 2:**
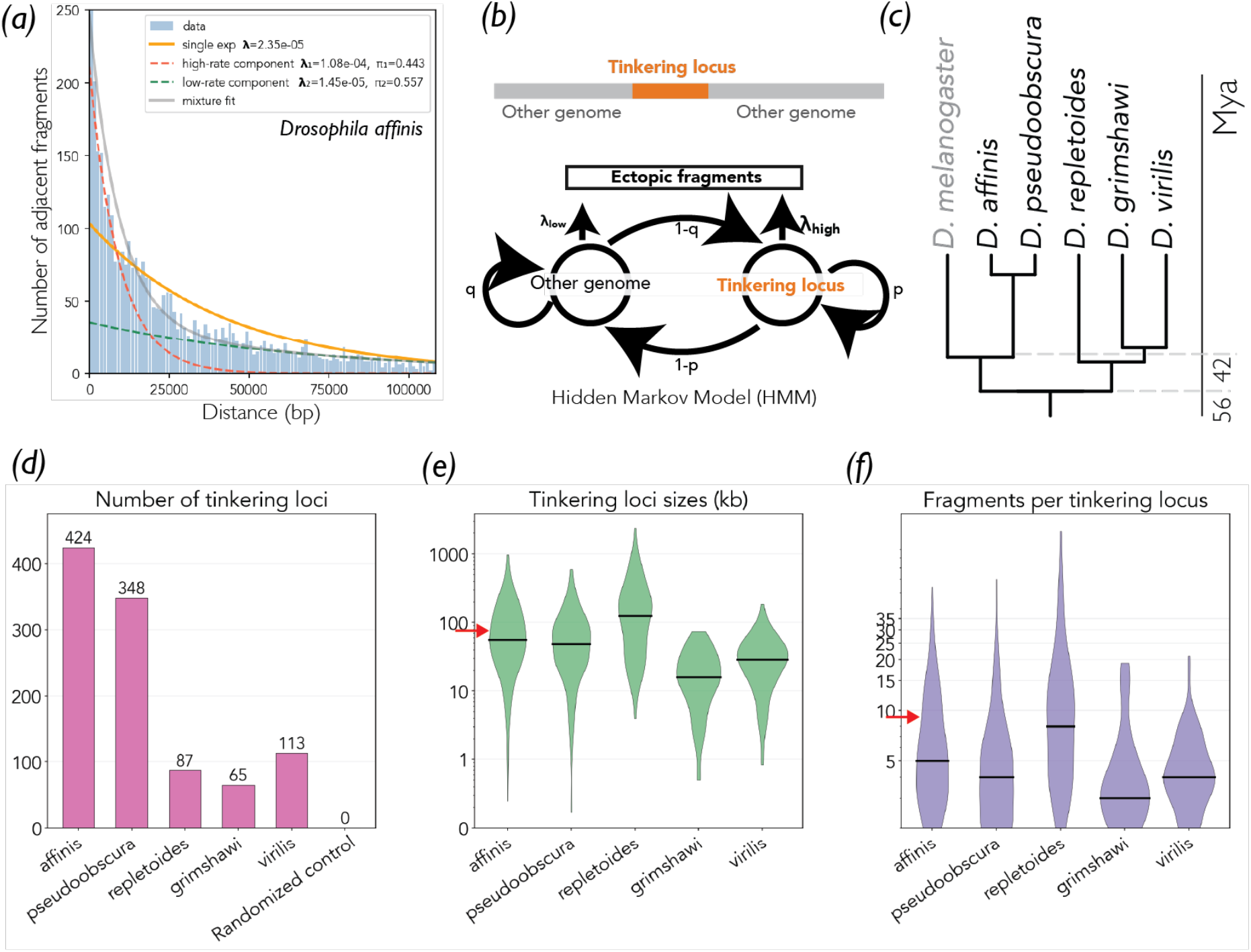
Tinkering loci are a general property of *Drosophila* genomes. (**A**): Fits of single (orange) vs. double (gray) exponential models to distances between adjacent dispersed fragments (blue bars) in *D. affinis.* There is an excess of fragments with close spacing relative to the single exponential, which the double exponential better captures. (**B**): Schematic of the HMM used to identify tinkering loci. The model has two states, tinkering loci and the rest of the genome, with different per-nucleotide probabilities of a fragment insertion, and different per-nucleotide probabilities of staying within the state or switching to the other. (**C**): Phylogeny of the *Drosophila* species analyzed with the HMM in D-F. (**D**): Number of tinkering loci found by the HMM in all species and a negative control genome with randomly shuffled fragment positions. (**E**): Size distributions of tinkering loci in all species. Horizontal black lines represent the median. Red arrow indicates the *goddard* duplicate locus (*affinis*) in Fig. 1B. (**F**): Distribution of number of fragments per tinkering locus in all species. Horizontal black lines represent the median. Red arrow indicates the *goddard* duplicate locus (*affinis*) in Fig. 1B.

Fragment spacings *λ*_1_,*λ*_2_ differed by ⪎10x in all species (Table S2), suggesting that a Hidden Markov Model (HMM) (Fig. 2B) may be able to use this signature to identify tinkering loci (Methods-7). I implemented such an HMM, which identified the examples from Fig. 1 in the *affinis* and *Chymomyza caudatula* genomes, and no loci in the negative control (Fig. 2D). This supports the visual intuition (Fig. 1) that these exemplar tinkering loci are indeed statistically anomalous compared to genomic background. I then applied the HMM to the five annotated *Drosophila* species. It identified many tinkering loci in each.

There is significant variation between species: *Drosophila affinis*, with the most (424), has more than 6x as many as *grimshawi*, with the least (65) (Fig. 2D). The sizes and numbers of fragments also vary between and within species (Fig. 2E,F). Examples of tinkering loci and background regions from *grimshawi* (visualized from the annotation files using IGV^48^) are shown in Fig. S2.

### Tinkering loci produce proto-genes at unusually high rates

The HMM identifies tinkering loci based on one of their core properties: an anomalously high concentration of ‘tinkering components.’ However, it does *not* consider how recently these components were acquired, or whether they are transcribed. These features are crucial to the capacity for rapid gene birth.

I chose *Drosophila affinis* for a closer analysis of tinkering loci on short evolutionary timescales. This species contains unusually many (Fig. 2D), and has four close relatives diverged by less than 8 million years (Fig. 1B), all with available “long-read” genome assemblies (Table S1), which minimize the possibility of assembly errors. I annotated these four species.

Long-read RNAseq (Methods-3) in *affinis* revealed that 56% of tinkering locus fragments are transcribed. This is below conserved genes (96%), as expected, and above unannotated control sequences (8%) and transposons (25%) (Methods-8; Table S3), indicating that fragments are read as biologically meaningful above background.

#### Abundant proto-gene birth at tinkering loci

Although most products of tinkering generally do not become conserved genes^49^, their creation is an essential step that provides the raw material for innovation. I therefore asked how quickly tinkering loci create new “proto-genes”^50^ – transcripts that encode new proteins, that have the potential to, like *goddard,* become functionally important – and what are their features. I identified groups of one or more fragments within tinkering loci that a) are transcribed and b) encode proteins that c) were created after *affinis* diverged from its closest relatives ∼3 million years ago^36^ (Methods-9).

I found 184 of these, each encoding one or more proteins in one or more of three categories (Fig. 3A, Table S4). 175 produce “singleton” transcripts, containing coding fragments from one protein. 41 produce “composite” transcripts, containing fragments from between 2 and 4 different proteins. Some of these transcripts encode multiple proteins (polycistrons^51^); others include fragments without coding capacity, e.g. lacking an intact open reading frame or in antisense orientation. These could nonetheless retain noncoding functions that are increasingly recognized to be common in coding sequence, like regulation of transcript stability or protein binding^52–54^. 4 produce “chimeric” transcripts, encoding a single protein derived from pieces of two different proteins. In total, these regions encode 234 proteins.

**Fig. 3:**
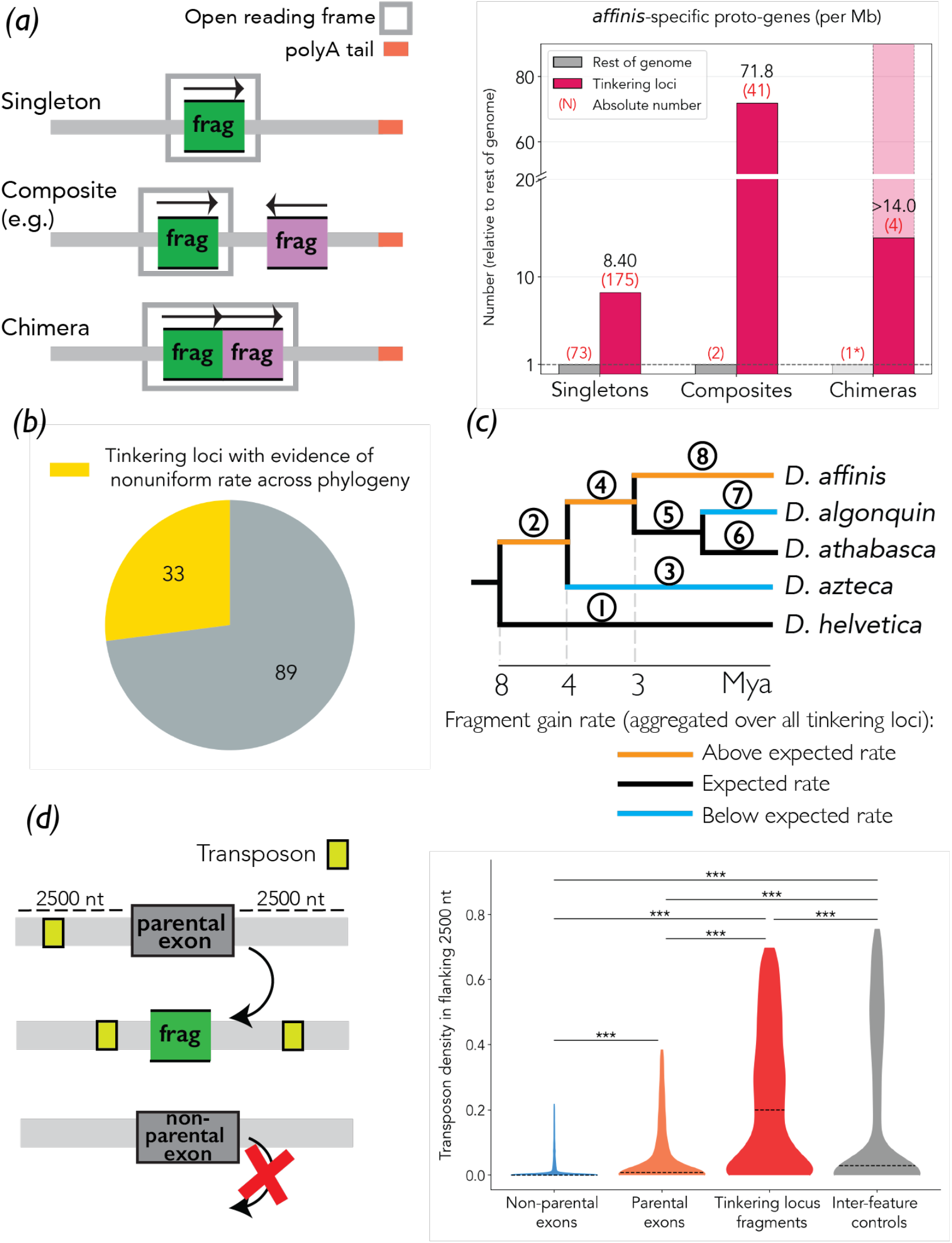
Products, evolutionary dynamics, and mechanisms of tinkering loci. (**A**): Left: cartoon schematic of categories of proto-genes as described in the text. Right: the number of *affinis*-specific proto-genes in each category created by tinkering loci, normalized per megabase to the number generated by the rest of the genome. Absolute numbers reported in the main text (red parentheses) are divided first by the size in megabases of the region type (tinkering loci / the rest of the genome), and then by the resulting value for the rest of the genome (gray), such that the gray bars are all normalized to 1. Shading on rightmost bar (chimeras) indicates the range of uncertainty resulting from 0 observed counts of chimeras outside of tinkering loci: assuming that the true average number is between 0 and 1, the solid bar indicates the bound of 1, and the translucent bar indicates the bound of 0. (**B**): Pie chart representing fraction of tinkering loci with evidence of heterogeneous fragment gain rate across the phylogeny (27%). (**C**): Inferred heterogeneity in the rate of fragment gain at *affinis* tinkering loci across the phylogeny. (**D**): Left: schematic of “parental” exons, which have produced a tinkering locus fragment by dispersed duplication; “non-parental” exons, which have not; and the definition of ‘flanking sequence’ (2500 nucleotides (nt) on either side of the coding sequence) used for analysis on right. Right: Transposon density (percent of sequence annotated as a transposon) in flanking sequences around parental exons, non-parental exons, tinkering locus fragments, and random control regions. Dashed black lines represent medians.

These fragments are derived from 143 ancestral genes (Table S4). Most appear at only one locus, but a few are at several. The latter include *ago2*, which functions in transposon/virus defense and whose extensive duplication under positive selection has been previously reported in *affinis*^55^, and *BTBD9*, which contains a *BTB/POZ* domain, a protein-protein interaction module involved in diverse cellular functions that has been noted for its adaptive evolvability^56^.

Fig. 4 shows an example tinkering locus with its *affinis*-specific chimeric proto-gene.

**Fig. 4:**
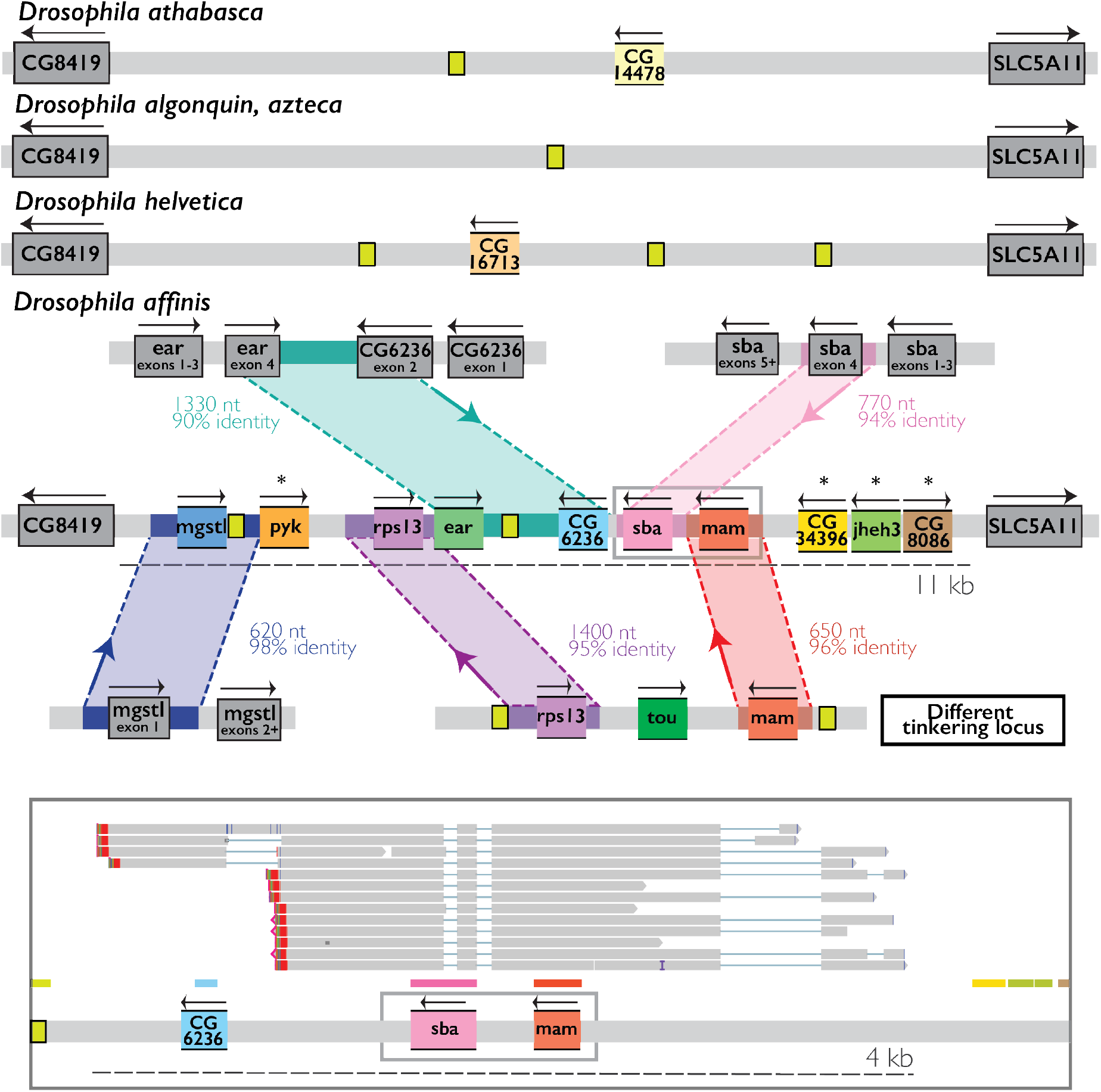
A tinkering locus generates an *affinis*-specific chimeric proto-gene. Legend for cartoons is the same as in Figure 1, with additional features as below. Top: the locus in outgroup species, with a low rate of fragment gain. Middle: the locus in *affinis,* and the inferred sources of the gained fragments. Ten fragments have dispersed to the locus from other regions of the genome since the divergence of *affinis.* Colors indicate sequences with high similarity, inferred to be recombination tracts. Starred fragments do not have flanking sequence matching parental exons, but are themselves partial internal segments of exons, such that such sequence is not expected. Direction of transfer is indicated by an arrow, inferred based on the presence of regions in outgroup species. Gray box represents the new chimeric ORF formed by the two enclosed fragments. Bottom: Iso-Seq data showing transcripts of three *affinis*-specific fragments, which together form an intact ORF with coding sequence derived from two ancestral genes, with the third fragment as a putative 3’ UTR on one isoform.

#### Tinkering loci produce proto-genes, especially composites, much faster than the rest of the genome

In the rest of the genome not annotated as tinkering loci, I found 73 regions producing proto-genes. All produce singleton transcripts; 2 also produce a composite transcript made from 2 ancestral genes; and none produce chimeric transcripts (Table S4).

Despite being a small fraction of the genome (Fig. S3), tinkering loci thus make more than twice as many proto-genes as all other regions, and more than 20 times as many composites. Normalized by the total amount of sequence that they comprise, tinkering loci create proto-genes overall 9x faster than the rest of the genome, and composite or chimeric transcripts 79 times faster (Fig. 3A).

#### Tinkering loci drive a higher proto-gene birth rate in *affinis*

Does *affinis,* given its unusually many tinkering loci, create new proto-genes faster than other species? Comparing these results with other studies’ is complicated by methodological heterogeneity. With this proviso and attempts to control for this heterogeneity (Methods-10), a study with similar methods^41^ suggests a 28x-higher rate of dispersed proto-gene birth in *affinis* than *melanogaster*, with an absolute lower bound of a 5x-higher rate overall, when only dispersed duplicates in *affinis* are compared to all duplicates in *melanogaster*. Another study^43^ suggests a rate of chimeric proto-gene birth in *affinis* 17 times higher than average between *melanogaster* and *Caenorhabditis elegans*. High gene birth rates have also been reported in *affinis* and closely related species *pseudoobscura* and *persimilis*^57,58^. It thus appears that tinkering loci significantly increase a species’ rate of proto-gene birth.

### Tinkering locus activity changes on short evolutionary timescales

I then asked: what are the rates and dynamics of tinkering at these loci?

The key property of tinkering loci is the accumulation of fragments. I first determined the “fragment flux” at each locus: the number of unique fragments whose presence is variable within the phylogeny, present in at least one species and absent in at least one other (Methods-11,12). In other words, this is the raw diversity in protein components that the locus has made available across the phylogeny. This distribution is long-tailed (mean of 4.2). Many loci have low or zero flux, while others are moderately to very active (Figure S4).

Low-flux loci could have a low but constant rate of fragment gain, such that observing no or few events within this phylogeny is expected. Alternatively, their activity could have changed over time. Variations in tinkering locus number within *Drosophila* (Fig. 2) and even the *affinis* group (Fig. S5) suggest that the latter can happen quickly. Are changes in tinkering locus activity observable within the *affinis* group? I assigned the gain of each fragment to a branch in the phylogeny and tested how many loci show changes in the rate of fragment gain across branches. This was the case for 27% of loci (p=10^-10^, Methods-12,13, Fig. 3B). Aggregating events across loci gave better power to quantify rate heterogeneity on individual branches (Methods-13). 5/8 branches deviated from the expected overall rate: 3/8 higher, and 2/8 lower, by 1.5-3x (Fig. 3C, Table S5). All branches traversed by *affinis* show elevated rates, an expected result, akin to a positive control, of the tinkering loci having been identified in *affinis*. The rest of the phylogeny shows idiosyncratic and frequent rate changes from branch to branch.

Tinkering loci also duplicate and rearrange fragments. I quantified “duplication flux,” the raw variation in duplicate copies of fragments, and duplication rate heterogeneity (Methods-11,12,13). Results are qualitatively similar to fragment gain (Table S5, Fig. S4), though duplication appears more evolutionarily labile, with larger flux extremes and more loci with rate heterogeneity (49%, p<10^-300^, Methods-13), as might be expected for a more common mutational process. 24% of loci showed at least one rearrangement within the phylogeny. A shared factor may drive the three tinkering modes: fragment and duplication flux are modestly correlated (r^2^=0.2, p=10^-5^; Methods-14; Fig. S4), and rearrangements predict meaningfully higher fragment (p=2*10^-7^, mean 6.4 vs 3) and duplication (p=6*10^-19^, mean 10.1 vs 2.3) flux (Methods-14).

As examples of the rate heterogeneity observed between tinkering loci, Figure 1B shows a locus with high flux and no rate heterogeneity, “active” over the whole phylogeny. By contrast, Figure 4 shows one with high flux and high rate heterogeneity (p=3*10^-5^; Methods-15), which has “turned on” only since the divergence of *affinis*.

### Tinkering uses ectopic recombination at transposons

What mechanisms enable the accumulation of fragments at tinkering loci?

I often noticed fragments lying within <2kb tracts of sequence matching their “parental” exons and extending into surrounding noncoding sequence (Fig. 4). This suggested ectopic recombination (ER), wherein a double-stranded DNA break is repaired using a non-allelic template locus, ‘mistakenly’ triggered by a region of shared sequence, as the underlying mechanism.

Consistent with ER, 49% of all tinkering locus fragments share ≥100 nt of flanking sequence (within 2500 nt, a scale typical of ER^59^) with their parental exon, versus a 3% baseline with random exons (Methods-16). Reverse transcription of the parental mRNA (retrotransposition) could also introduce shared sequence flanking exons. I found that 7x more fragments share flanking sequence with the parental exon at the DNA level, but not at the RNA level, than vice versa (Methods-16). This suggests ER over retrotransposition as the dominant mechanism.

I also noticed that transposons appeared common at tinkering loci, often juxtaposing dispersed fragments (Figs. 1, 4). This suggested transposon sequences, repeated in many places around the genome, as an important sequence substrate for tinkering locus ER. Recombination at transposons is a known mechanism for gene birth by tinkering outside of tinkering loci^60^.

Annotated transposons comprise significantly more of flanking sequence around “parental” exons that have given rise to tinkering locus fragments than around exons that have not (Fig. 3D, median 0.01 vs. 0, Mann-Whitney U p=10^-92^, Methods-17), consistent with involvement in their dispersal. They are also denser around tinkering locus fragments than non-exon, non-fragment control sequences, which should be minimally functionally constrained and thus approximate neutral background levels (Fig. 3D, median 6x, Mann-Whitney U p=10^-56^, Methods-17). Enrichment does not appear specific to a single transposon family or even broad class: it includes sequences with signatures characteristic of DNA transposons, both classical and Helitrons, and retrotransposons, both LTR and non-LTR (Methods-17,4-4b, Table S6). This is consistent with the role of transposons primarily as widely distributed shared sequences that enable gene fragment spread by ER.

### Tinkering loci form networks that enable their products to duplicate and spread

I noticed repeated *affinis*-specific duplication of a chimeric gene whose coding sequence is comprised of fragments of *Rcc1* and *Txl*. Because recurrent duplication is suggestive of function, I analyzed its properties and evolutionary history (Methods-18, Fig. 5).

**Fig. 5:**
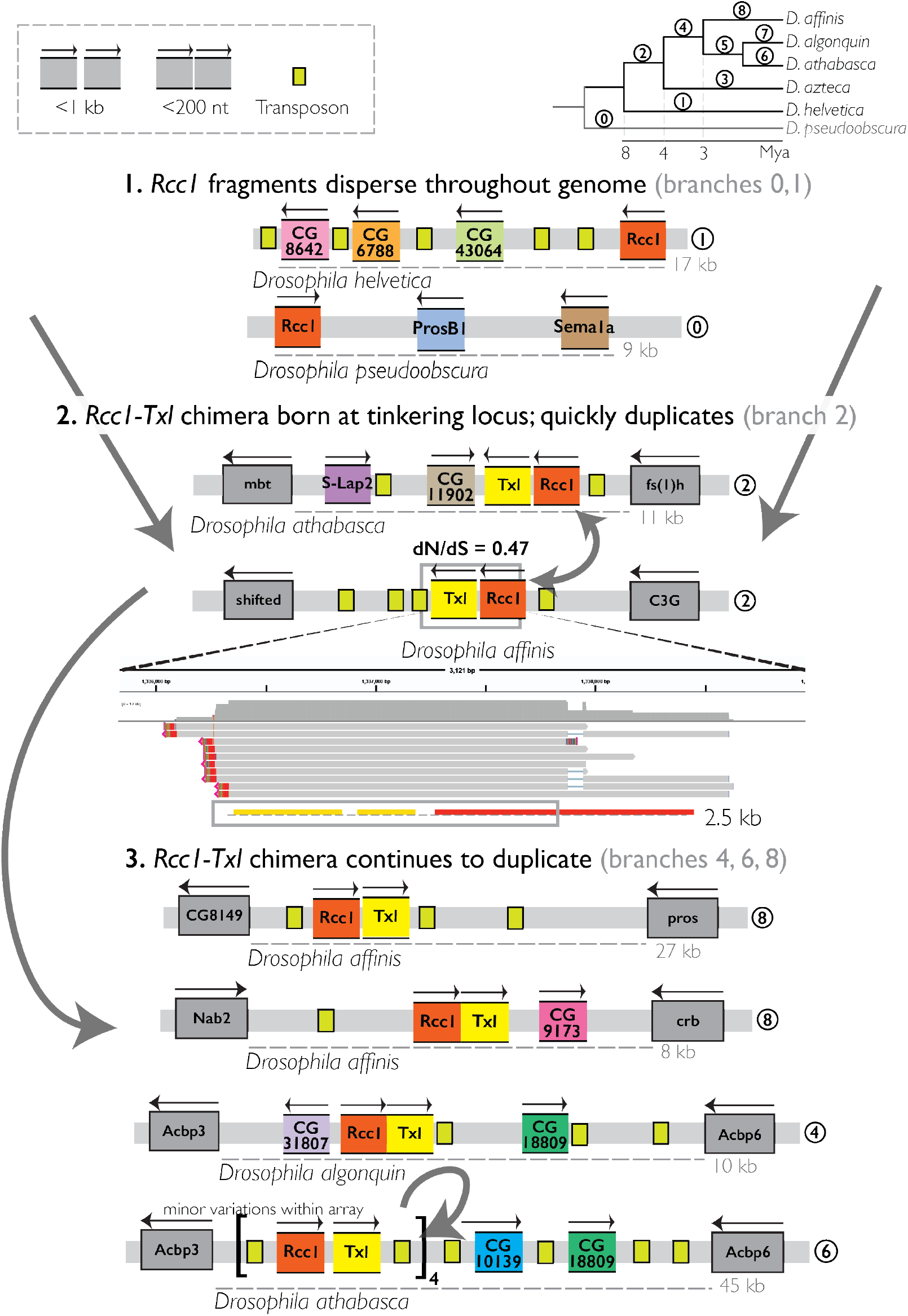
Creation and spread of a chimeric gene via tinkering loci. A schematic illustration of inferred steps in the creation and evolution of a *Rcc1-Txl* chimeric gene via *affinis* group tinkering loci, as described in Methods-18. Legend is the same as for Figures 1 and 4; arrows represent duplication of individual fragments (Step 2) or the new chimera (Step 3). The chimera formed from the duplication of fragments from *Txl* and *Rcc1* to one of the two loci shown in Step 2 and subsequently spread to many other loci in the group. Circled numbers next to loci indicate the inferred branch, corresponding to the phylogeny in the top right, of the depicted event. The order and direction of transfer of events within Step 2, and the precise source of each duplicate copy in Step 3 (except the bottom locus, where tandem duplication, due to its mutational accessibility, is assumed), were not inferred; arrows are meant to represent this ambiguity. For ease of illustration, details of the depicted loci are shown only for the indicated species, even when the chimera exists in multiple (fragments at tinkering loci often vary between species).

I find that this chimera was created recently, within the *affinis* group, and that the full evolutionary history around its formation and spread is complex. First, *Rcc1* fragments began to proliferate in the genome well before formation of the chimera: in all *affinis* group species, as well as outgroup *pseudoobscura*, fragments are present in many copies, including many at tinkering loci. The chimera was then born at a tinkering locus ∼4 Mya, in the common ancestor of *affinis, athabasca, algonquin,* and *azteca.* Soon after, it duplicated to a second locus. One of these two copies degraded in two species; the other was retained in all four species, and has a dN/dS of 0.43, indicative of purifying selection. The chimera then duplicated four more times, including several specifically in *affinis,* mostly to other tinkering loci. The origin at and duplication to tinkering loci parallels the history of *goddard*.

## Discussion

Tinkering loci are distinct regions of *Drosophila* genomes, distinguishable from background quantitatively and by eye, that meaningfully accelerate the rate of gene birth by “tinkering.” They do this by accumulating and re-transcribing unusually high numbers of dispersed gene fragments. To summarize, they appear to have four basic properties. First, they produce new “proto-genes” much faster than the rest of the genome. As in other cases of gene birth, most will likely not acquire functionality, but an increased rate correspondingly increases the chance of hitting upon those that do, like *goddard* and the *Rcc1-Txl* chimera. Second, this acceleration is especially large for “composite” transcripts, which combine existing gene pieces in new ways. This allows efficient exploration of the combinatorially large space of new functions that can be generated by combining existing ones, which is usually an unlikely outcome in large animal genomes. (Even when composite transcripts do not form an intact protein, they may still have novel functions, endowed by noncoding functions of protein-coding sequences^52–54.)^ Third, tinkering locus activity is highly dynamic, changing on short evolutionary timescales. Fourth, their causal mutations are consistent with ectopic recombination, a universal eukaryotic process, acting on sequences of all major classes of transposons, which are abundant in animal genomes.

Tinkering loci are unusual because of their deviation from the standard view in which tinkering occurs via mutations that are more or less genomically unpatterned. By contrast, these are highly nonrandom spatial patterns of mutations, with meaningful consequences for the rates and outcomes of molecular innovation. I see them of as importance for two reasons rooted in this feature. First, they represent a qualitatively new strategy for increasing the rate of gene birth. Their evolutionary dynamism suggests that this strategy may contribute to variation in the rate of molecular and thus phenotypic innovation – the “tempo”^61^of evolution – across taxa and over time. Second, between their determinants of transposon sequences, which appear to direct recombination-driven duplication, and possible additional enabling factors, like patterns of DNA damage (to trigger recombination), chromatin accessibility, or 3D architecture (to make donor and recipient loci accessible and bring them into contact), they suggest that nonrandom patterns of “tinkering mutations,” from which they arise, *can be encoded by the genome itself*.

The second point has several broader implications. First, if these patterns are encoded by the genome, then the variation within them can be selected upon, either due to direct effects of their genomic determinants (e.g. fitness effects of transposon insertions) or as linked to the molecular innovations that they produce. The high variability in genomic transposon content within populations^62^ and over time^63^ may make many variants available neutrally, giving selection many possible substrates. In principle, this seems an accessible and mechanistically plausible route to the evolution of “evolvability.”

Second, if the genome encodes nonrandom mutational patterns, it seems possible that these patterns could take forms other than single obviously anomalous loci. Beyond their obvious density of duplicates, tinkering loci exhibit a behavior characteristic of hubs in networks: they do not just receive duplicates, but disperse them -- including both their products (*goddard, rcc1-Txl*) and individual fragments (Fig. 4, in which a piece of the new chimera comes from another tinkering locus rather than directly from the ancestral gene) – to other loci, including and especially fellow hubs. I speculate that this is not just a useful metaphor: that tinkering loci are hubs in genome-wide “recombinational networks,” in which nodes are loci and edges connect loci whose shared sequences place them in “recombinational contact.” This network defines the structure upon which tinkering and its resulting molecular innovations play out. It may include other nodes with nonrandom properties different from those of tinkering loci; primate segmental duplications^64,65^ appear to be an example. Its structure may also be highly nonrandom in ways that are apparent not from inspecting individual loci but only from observing the network as a whole. Its products may bear the fingerprints of this structure, as in the more straightforward somatic analogs of V(D)J recombination^66^ and ciliate genome rearrangement ^67^. An understanding of the network may thus give insight into cryptic structure in its products. These products may include not just the protein coding components used here, but also regulatory elements, which are increasingly recognized to be made of many modular components^68^, and which underlie intricate gene regulatory networks that are key to animal complexity^69^. In animal genomes, recombinational networks are likely large, densely connected, and dynamic, changing with transposition events and duplications themselves. Networks with these properties can give rise to complex, nonlinear, and unintuitive phenomena. This way of thinking thus suggests a speculative framework that may also shed light on many features of the *mode*^61^ of evolution in animals.

Tinkering loci were discovered while investigating the origin of *goddard*, an essential new animal gene hypothesized to have originated de novo based on a lack of detectable homology to other genes^32^. Because fragments at tinkering loci are often initially small and subsequently disrupted by structural mutations, detectable similarity to the ancestral protein is expected to erode rapidly^19^. Tinkering loci may thus be a source of many other genes currently believed to be de novo.

## Acknowledgements

I am grateful to Joshua Akey, Michael Levine, Julien Ayroles, Andrew Murray, and Sean Eddy for mentorship and useful conversations; Joshua Akey for laboratory space and computational resources; Sean Eddy for computational resources; Dayna Akey, Michael Levine, Julien Ayroles, Julie Peng, Emmanuel D’Agostino, and members of the Levine and Ayroles laboratories for experimental support and resources; Wei Wang and Jean Arly Volmar for their experimental work through the Princeton Genomics Core; Joshua Akey, Andrew Murray, Michael Levine, Christopher Catalano, Pavan Choppakatla, and Thomas Tullius for feedback on the manuscript; and the Lewis-Sigler Scholars program at the Princeton Lewis-Sigler Institute for Integrative Genomics for financial support and scientific independence.

## Declaration of interests

The authors declare no competing interests.

## Methods

Unless otherwise specified, analyses use custom Python and/or bash scripts, available at https://github.com/caraweisman/Weisman_tinkering_loci_2026.

### 1. Identification of *goddard* orthologs and loci

In *melanogaster* and other *Drosophila* species, *goddard* lies in an intron of conserved gene *omega* ^32^. According to the r6.68 release of FlyBase, the exons between which it lies are exon 2 (coding) and exon 3 (noncoding). Exons 3 and 4 (which is very short) are not well-conserved in other *Drosophila*, so for the purpose of searching for orthologs in outgroups, I define the locus as the region between exon 2 and the nearest coding exon (5), coordinates (with a small amount of flanking sequence on either side) 3L:14679497..14740596 (61.1 Kb).

In the *Drosophilidae* assemblies listed in Table S1, I used TBLASTN to identify exons 2 and 5 of *omega,* as annotated in FlyBase release FB2023_05. I then performed TBLASTN and phmmer searches for the *goddard* coding sequence in this region in an iterative manner: I began with the *melanogaster* sequence studied experimentally in previous work ^32–34^, searched for it in the identified orthologous loci, and, if a hit was returned at E<0.001, manually inspected the putative ortholog’s sequence and alignment to exclude low-quality hits due e.g. to repetitive sequence. I then repeated the process using newly identified orthologs as query sequences. This enabled stepwise detection of *goddard* orthologs in species increasingly divergent from *melanogaster*, softening the difficulty that *goddard*’s fast evolving sequence poses for homology search ^32^. I identified orthologs in *Scaptodrosophila inornata*, *lebanonensis,* and *latifasciaformis*, indicating that *goddard* was present in the common ancestor of *Drosophila* and *Scaptodrosophila*.

### 2. Expected number of family-specific genes at *goddard* locus

In addition to *goddard*, *CG13476* and *CG13471* appear to be specific to *Drosophilidae.* I used data from ^19^ which performed a homology search for all *melanogaster* proteins in 11 insect outgroups, including four non-*Drosophila* Dipterans *(Musca domestica, Ceratitis capitata, Anopheles gambiae, Aedes aegypti)*, to determine the number of *melanogaster* genes with no detectable homologs in species outside of *Drosophila* (1100). I then performed a binomial p-value calculation to calculate the probability of 3 of these 1100 genes lying in a 60kb region by chance, corresponding to k=3 successes, p(success) = 1100/180,000,000 (number of lineage-specific genes per position in the genome), and N=61,000 (number of positions in the locus, per <u>Identification of goddard orthologs</u> above).

### 3. Drosophila affinis Iso-Seq

I purchased *Drosophila affinis* strain 14012-0141.00 from the National Drosophila Species Stock Center (NDSSC). This is the strain used for the assembly analyzed here, from Kim et al. 2024 ^70^. (I selected *affinis* as the focal species because it was the only readily available live strain in the *affinis* group.)

I sorted male and female flies, made simple by the distinctive red coloring of *affinis* group male testes, visible through the abdomen. Approximately 40 male and 40 female flies, ages 3-5 days, were anesthetized with CO_2_, frozen on dry ice, stored at −72°C, and mechanically homogenized with a pestle in Zymo DNA/RNAShield. RNA was extracted using the ZymoBIOMICS Quick-RNA miniprep kit (Zymo Research, R1655) and eluted into 30 uL. Iso-Seq libraries were created according to the ‘Preparing Iso-Seq® v2 libraries’ protocol from PacBio, with barcoded cDNA primers for multiplexing with three other *Drosophila* samples. Libraries were pooled, bound, and cleaned according to the Revio SPRQ™ Polymerase kit. Libraries were then loaded onto the Revio instrument and run using the application presets. This resulted in 5.1-5.3M reads, each corresponding to a transcript, per sex. For all analyses here, I pooled reads from both sexes, for a total of 10.4M reads.

### 4. Genome annotation

#### Rationale

Existing eukaryotic gene annotation pipelines, optimized for identifying high-confidence genes, are by design poorly suited to the quite different task of identifying gene fragments, enriched for features (short length, fragmented ORFs) unfavorable to this end. Similarly, much pseudogene identification software aims to find full-length genes pseudogenized in situ by an inactivating mutation ^47^. A custom approach to identify gene fragments was therefore necessary.

The desired output from an annotation pipeline to study tinkering loci will both identify gene fragments and determine the ancestral genes from which they are derived, so that the underlying molecular evolutionary mechanisms and biological features of products can be studied. Additionally, a comparative analysis based on these annotations also requires that they be straightforwardly comparable across species. The simplest and most conservative way to achieve both aims is to annotate the same conserved core of ancestral genes, in which there is high confidence and available functional information, and gene fragments derived from them, in all species.

I took this approach in developing a custom pipeline to annotate conserved genes, gene duplicates, and gene fragments. I used the *Drosophila melanogaster* protein annotation (FlyBase Release 6.54, via NCBI GCF_000001215.4) as the reference protein set, motivated by its being the highest-quality Dipteran annotation with the most experimental information. This is a rough proxy for the protein repertoire in the common ancestor of *melanogaster* and the othe*r Drosophila* studied here.

#### Limitations

This approach has the limitation that lineage-specific genes (with no detectable homologs in *melanogaster*) and fragments derived from them will be missed; the numbers of new genes reported here are therefore likely an underestimate. However, the relatively short evolutionary timescales involved in this analysis mean that the number of such genes is unlikely to be substantial ^19^, unless lineage-specific genes are themselves likelier to be duplicated at and by tinkering loci.

The detectability of a species’ homologs of *melanogaster* proteins is known to depend on evolutionary distance ^19^, meaning that results from this approach will only be comparable between species at similar evolutionary distances from *melanogaster.* All species annotated with the pipeline in this study roughly satisfy this criterion (Fig. 2) ^36,71^, with the exception of *Chymomyza caudatula*, which was annotated only as a positive control for the tinkering locus HMM, and was not included in comparative analyses. Further confidence that the evolutionary distances of the species in question are not the primary determinant of the tinkering locus analysis results is provided by the finding that the differences in tinkering locus content within the *affinis* group, with the same divergence time from *melanogaster,* are comparable to those between the other species, which have slightly different divergence times (Fig. S5, Fig. 2C).

## Method

At a high level, the pipeline first identifies homologs of each *melanogaster* protein through sequence similarity. It then considers additional sequences in the genome that also have significant sequence similarity to these proteins, and classifies them as gene fragments or duplicates. It also identifies transposon sequences.

The accessions for the RefSeq/Genbank genome assemblies annotated with this pipeline are listed in Table S1.

A more detailed summary of the annotation methodology is below. The resulting annotations, as GFFs compatible with the Integrated Genomics Viewer ^48^, are available at https://github.com/caraweisman/Weisman_tinkering_loci_2026.

### 1. Homology search of *Drosophila* proteins

The annotation pipeline first performs a TBLASTN search of the RefSeq *melanogaster* protein annotation against the genome assembly of the target species with an E-value cutoff of 0.01, meant to strike a balance between allowing detection of short protein fragments, which due to their length will have relatively high E-values, and controlling the false positive rate.

### 2a. Conserved homolog identification

The pipeline first uses these results to identify a conserved homolog of each *melanogaster* protein in the target genome. This iterative process begins with an “anchor” hit, chosen by E-value in ascending order (beginning with the lowest and most significant). It then attempts to expand outward from the anchor hit’s location on the target genome to find other adjacent hits within 50kb (well above the length of the average intron) whose strand, orientation, and position in the *melanogaster* protein and target genome are consistent with it being another exon of the protein (i.e. fusions and fissions of exons are not penalized; exons must merely come in the same order as they do in the *melanogaster* protein). This extension terminates when another such segment cannot be found. If the total portion of the *melanogaster* protein covered by all hits in this extension group does not exceed 60% of its length, the process repeats with the next anchor hit, and terminates with a hit that does exceed the threshold. If no anchor hit results in a set of exons that exceeds the threshold, the first anchor hit, with the lowest E-value, and its group are chosen as the homolog. Because E-value is itself strongly correlated with length, this process is intended to prioritize longer hits to *melanogaster* proteins, less consistent with being a gene fragment. These parameters were optimized empirically, through visual inspection of results and benchmarking analyses. This process always finds at least one homolog for each *melanogaster* protein if it has any significant hit in the target genome. This is an intentionally conservative choice which makes it impossible to call protein fragments unless the gene has more than one significant hit in a genome. This conservatism, which risks including fragments or pseudogenes as orthologs in favor of deflating the false positive fragment rate, is also another way in which the goals of this approach differ from those of traditional eukaryotic annotation approaches, and was inspired by unsatisfactory early pilot experiments with such methods.

### 2b. Homolog identification benchmarking

On average, 96.2% of *melanogaster* proteins had a homolog successfully identified in each of the five species annotated in Figure 2, with an average of 86.8% of the *melanogaster* length included in the annotation (counting each position in the *melanogaster* protein at most once, even if e.g. an exon is duplicated):

| Species | % *melanogaster* genes annotated | Average % *melanogaster* gene length included in annotation |
| --- | --- | --- |
| *Drosophila affinis* | 95.9% | 85.8% |
| *Drosophila repletoidea* | 96.8% | 88.3% |
| *Drosophila pseudoobscura* | 96.9% | 88.4% |
| *Drosophila grimshawi* | 95.4% | 85.4% |
| *Drosophila virilis* | 96.0% | 86.1% |

These numbers are similar to those reported upon the sequencing of the *pseudoobscura* genome ^72^ (>90% of *melanogaster* genes with homologs detectable by TBLASTN, covering 93% of the *melanogaster* gene length).

I also find that 96% of genes annotated as homologous are expressed in *Drosophila affinis* (main text), compared to 56% of annotated tinkering locus fragments, 25% of transposons, and 8% of length-matched intergenic control sequences. This much higher expression than all other categories supports the annotation of these genes as conserved.

### 3. Homology search for bacterial genes

The pipeline also seeks to identify genes horizontally transferred from bacteria, a known source of novel *Drosophila* genes ^73,74^. As the first step, the pipeline performs a TBLASTN search with an E-value cutoff of 0.01 of three bacterial annotations against the target species’ genome. These bacteria are *Acetobacter pasteurianus* (RefSeq accession GCF_009914215.2) and *Lactobacillus plantarum* (RefSeq accession GCF_009913655.1), known colonists of the *Drosophila* intestine ^75^, and *Wolbachia*, a known *Drosophila* endosymbiont (RefSeq accession GCF_016584425.1).

### 4. Homology search for transposons

The pipeline also seeks to identify transposons.

### 4a. DFAM-based search

As one step, the pipeline performs a BLASTN search with an E-value cutoff of 0.01 of all 881 curated Dipteran transposon sequences from the database DFAM ^76^ against the target species’ genome.

### 4b. HMMER-based search

Because transposons can evolve quickly and be horizontally transferred, nucleotide-based similarity searches are often insufficient to identify them. This is of meaningful concern because the above Dipteran DFAM sequences are entirely taken from either the *melanogaster* group of *Drosophila* or from mosquitoes, both more than 30 My diverged from the *affinis* group, making failure of nucleotide homology search more likely. Qualitative indications of this problem came upon running an earlier version of the pipeline without the present step and noticing many transcribed regions that themselves had no transposon annotation, but were surrounded by transposon annotations, suggestive of being an unannotated transposon within a transposon hotspot. A more sensitive search is therefore desirable. I curated a set of 13 PFAM ^77^ domains known to be associated with transposons:

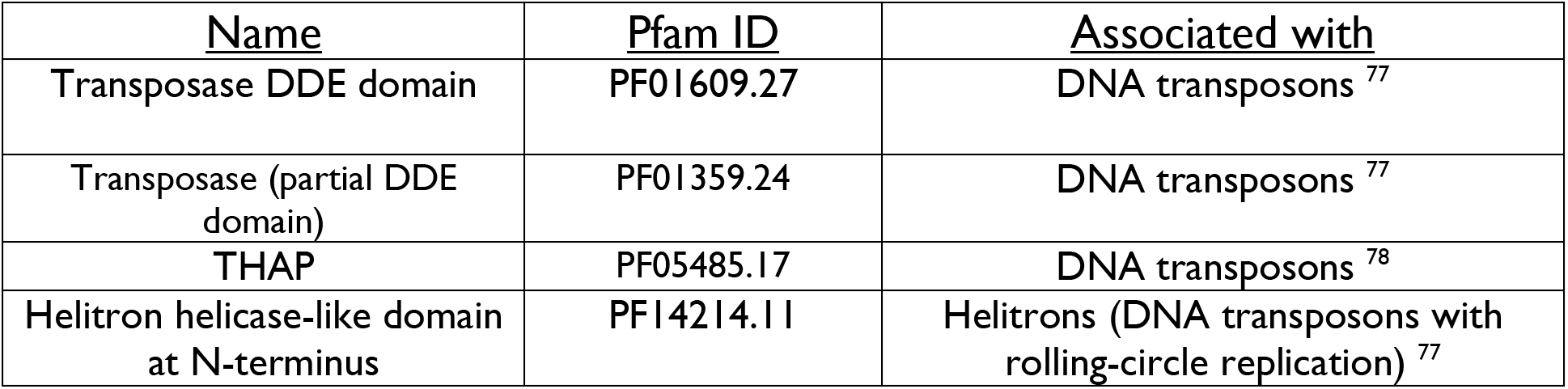
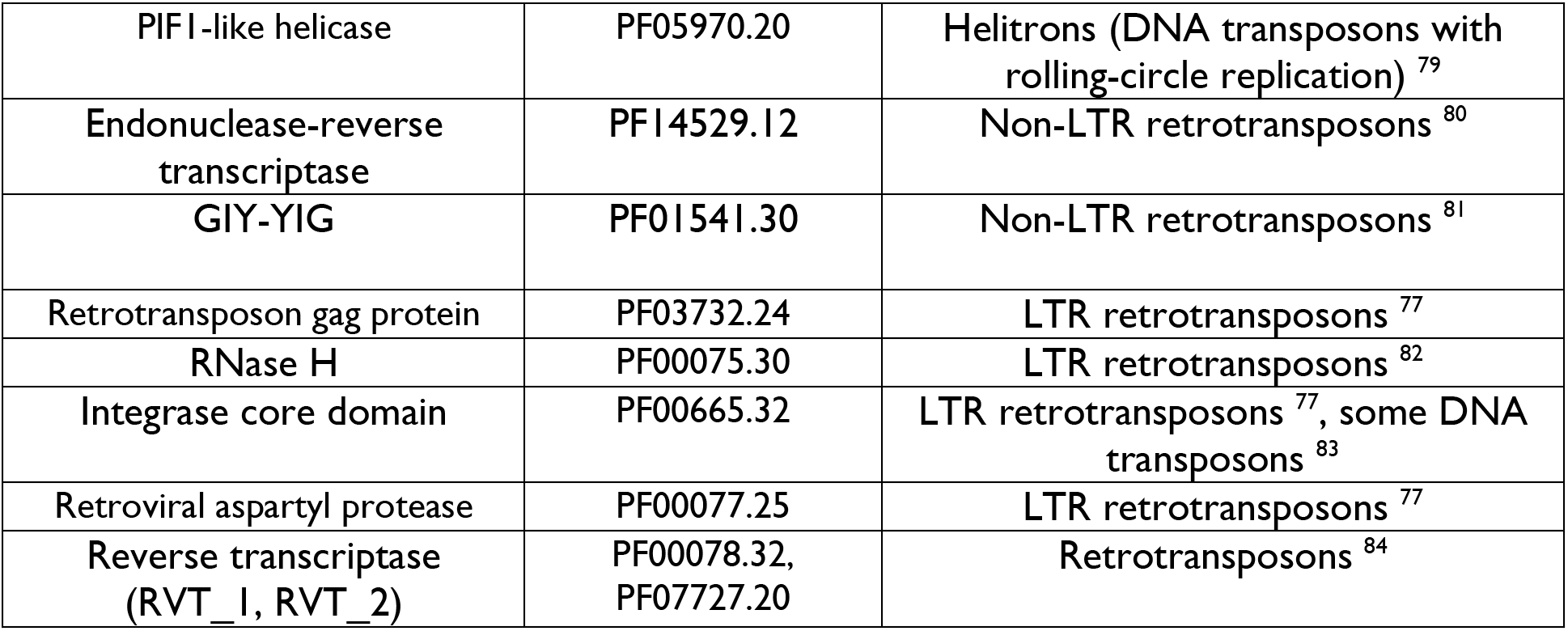

I then performed a six-frame translation of the target genome using the HMMER ^85^ utility esl-translate and performed a HMMER hmmscan search of these domains against the genome, with an E-value cutoff of 0.0001.

### 5. Gene duplicate/fragment, transposon, and bacterial horizontally transferred gene identification

Next, the pipeline jointly uses results from homology searches described above: a) hits to *melanogaster* proteins that were not called as orthologs, which can be thought of as “extra” hits, left over after all orthologs have been accounted for to the extent reasonably possible; b) hits to the bacterial protein annotations described above; and c) hits to the transposon databases described above. All hits in these three categories that overlap (≥50 bp) with regions already annotated as conserved orthologs are removed, prioritizing these conserved gene annotations. The remaining hits are kept if a) they do not overlap with any other remaining hit, or if b) they overlap with another remaining hit but have the lower E-value. Hits to *melanogaster* proteins resulting from this step are considered fragments/duplicates; hits to bacterial genes are considered putative horizontally transferred genes (very few of these result, and I do not discuss them here); hits to transposons are considered transposons.

### 6. Parsimony-based homogenization of gene fragments

I found that the above process can result in minor differences in E-value causing nearby fragments of what is likely the same *melanogaster* protein (e.g. resulting from the duplication of multiple exons, fragmentation of single exons after duplication, or subsequent local tinkering like duplication/rearrangement following the dispersed duplication event) to be annotated as derived from different, but related, paralogous genes. The likelier correct annotation, and an interpretation that enables the most conservative downstream estimates of the underlying number of distinct dispersed duplication *events,* is to assume, where possible, that neighboring fragments are derived from the same ancestral gene. To achieve this, I implemented a step in which the annotations of plausibly paralogous nearby hits are “homogenized,” made to match a single common ancestral protein. I first performed an all-by-all BLASTP search on the *melanogaster* proteome with a permissive E-value cutoff of 0.01, intended to include even remote paralogs and be maximally conservative in inferring distinct duplication events. I then re-analyzed the list of gene fragments annotated in above step (5) by determining whether any two genomically adjacent fragments within 20kb fall into the same cluster. If so, I selected the hit with the lower E-value; determined whether the neighboring hit(s) are included in the cluster of its annotation; if so, checked the raw TBLASTN search results to determine whether these neighboring hits themselves overlapped by more than 50 bp with a hit to that hit’s annotation; and, if so, replaced the current hit by that hit, thereby switching the annotation of neighboring fragments to those that, parsimoniously, come from the same protein.

### 7. *Drosophila affinis* only: RNA-based profile HMM annotation

Specifically for *Drosophila affinis*, the only species for which there was available long-read RNA-seq data (generated here, above), I also performed a third transposon annotation approach. I repeated the PFAM transposon domain hmmscan described above on unique transcripts (found by clustering reads using MMseqs2 ^86^, --min-seq-id 1 -c 1 --cov-mode 0) that, when mapped to the genome, did not overlap any conserved *melanogaster* genes or any gene fragments or duplicates. The resulting transcripts were considered putative transposons. I then clustered these transcripts (aggregated from both sexes) with MMseqs2 ^86^ easy-linclust --min-seq-id 0.9 -c 0.9 --cov-mode 0. Finally, I performed a BLASTN search with an E-value cutoff of 0.01 of these clustered sequences against the *affinis* genome to identify other copies of these transposons that are not actively expressed (e.g. degraded copies).

The rationale for this additional transposon annotation step was threefold. First, reducing the search space for transposons from the entire genome to expressed sequences increases power significantly, with a biologically compelling basis: active transposons are expected to be transcribed. Second, this allows identification and annotation of much more of the transposon sequence, including regions other than canonical transposon domains, which may be especially likely to have diverged from identifiable DFAM sequences and not included in domain-based searches. Third, transposons with active expression are likelier on balance to be biologically important, improving downstream analyses.

Because long-read RNAseq data was only available for *affinis*, I kept this annotation as a separate file and did not integrate it, hoping to preserve commensurability between different species’ annotations. It is used for the transposon analysis in the main text and is available as a separate file in the supplementary information for that section.

### 5. Inference of number of dispersed gene duplicates/fragments in *Drosophila* <u>genomes</u>

Counting distinct gene fragment or duplicate features individually as they are listed in the above annotation file (akin to the way that individual exons are listed: as separate features) is not appropriate, as it will separately count e.g. a) adjacent exons of the parental gene, resulting from the same dispersal event, or b) adjacent fragments within a single exon, generated by small deletions within the exon after dispersal, not of interest here. These separate counts do not reflect bona fide tinkering activity like independent dispersal, local duplication following dispersal, or local rearrangement following dispersal.

To conservatively avoid such overcounting and focus on counting segments separately only when they reflect tinkering mutations of interest, I took the above annotations, extracted gene fragments, and merged into a single entry adjacent fragments that are within 10 kb, are annotated as the same protein or in the same cluster (Methods-4.6), are on the same strand, and leave a gap of less than 200 amino acids relative to their parental protein. (Note that the genome annotation process, Methods-4, partially facilitates this by homogenizing the annotations of neighboring fragments with these properties, but does not yet merge their entries, for the same reason that different exons are listed separately in standard annotations: it is useful to preserve information about their location on the genome. This step formally merges these entries into one so that they can be counted together, but is only used for this purpose.) These features were chosen to select cases in which the hypothesis of a single dispersal event with no meaningful subsequent tinkering is, conservatively, likelier than that of bona fide tinkering. The merged fragments were counted once each.

To identify the number of dispersal events, the results of tandem duplication must be excluded. I thus excluded merged fragments for which either of the closest two “flanking” conserved orthologs of a *melanogaster* protein, one on either side, matched the fragment’s annotation, or the annotation of another gene showing detectable homology to it (in its cluster, Methods-4.6).

I also removed two classes of genes (histones and NUMTs, mitochondrial gene derivatives) that form many arrays throughout the genome, inflating dispersed fragment counts and downstream analyses in ways misleading as to the overall dynamics.

### 6. Single vs. double exponential model fit to nearest-neighbor distances of dispersed fragments

I first used the above list of dispersed fragments (merged, to reflect the number of discrete events) to calculate the distance between each adjacent fragment pair. The first and last fragments on a contig only had one pairwise distance; all others had two. These distances form the genome-wide distribution of nearest-neighbor fragment distances.

To assess the relative fit to the two models (the null model, a single exponential *e*^−*λ*x^, representing a single average distance 1/*λ* between nearest-neighbor fragments in the whole genome, and the alternative model, the sum of two exponentials, *π*_1_*e*^−*λ*1x^ + *π*_2_*e*^−*λ*2x^, where *λ*_1_,*λ*_2_ are the inverses of the mean spacings and (*π*_1_+ *π*_2_ =1), representing two types of regions in the genome with distinct average distances, the shorter distance corresponding to tinkering loci) of this distribution of nearest-neighbor fragment distances, I first fit the data to the models by estimating the maximum likelihood parameters for each model (*λ* for the null model; *λ*_1_,*λ*_2_, and *π*_1_ for the alternative model). I did this by optimizing the negative log likelihoods of each model using the Nelder-Mead simplex algorithm as implemented in Python scipy.optimize.minimize ^87^, stopping at convergence, defined at 10^-10^ for both the log likelihood and parameters, or at a max of 50,000 iterations. The *λ* parameters were optimized in log space to enforce positivity, and *π*_1_ was constrained to the range (0,1) by a large penalty term. The Nelder-Mead initialization point for the null model was the empirical inverse mean spacing calculated directly from the full empirical distribution (the maximum likelihood estimator). For the alternative model, with its potentially complex likelihood surface, I used a half-grid of initialization points: *π*_1_ ∈ {0.05,0.1,0.2}, *λ*_1_/*λ*^0^ ∈ {5,20,50}, and *λ*_2_/*λ*^0^ ∈ {0.1,0.3,0.5}, where for consistency *λ*_1_>*λ*_2_, i.e. always making the first component the tinkering locus component. This grid was intended to reflect biologically plausible and conservative assumptions about tinkering loci (that they contribute overall a minority of fragments, that the average spacing of fragments in tinkering loci is no more than 50x denser than the overall average, that the average spacing outside of tinkering loci is no sparser than 10% of the overall average). It is worth noting that failure to fully optimize the parameters in the alternative model would favor the null model, the opposite of what is observed here, and that the ranges here serve merely as initialization points, which the optimizer can go beyond, as in Fig. 2 / Table S2.

After finding these best fit parameters and their corresponding log likelihoods for both models, I compared the models via three criteria.

<u>1. The Akaike Information Criterion</u>, AIC = 2k-2l, where k = the number of free parameters (3 for the alternative model, 1 for the null model) and l = the log likelihood. Lower values indicate the preferred model.
<u>2. The Bayesian Information Criterion</u>, BIC=k*log(n) – 2l, where values are the same as above, and n = sample size. Again, lower values indicate the preferred model.
<u>3. Likelihood Ratio Test p-value.</u> I performed a likelihood ratio test (LRT) by computing the likelihood ratio statistic 2(l_alternative_– l_null_), and then generating an empirical null distribution by generating 1000 samples from a null model with *λ* equal to the average value in the dataset, fit both models to this sample, and computing the likelihood ratio statistic. The p-value of the likelihood statistic was calculated empirically, as the fraction of simulated replicates where the likelihood statistic equaled or exceeded the observed statistic.

As a negative control for this workflow, I produced a dataset of fragment distances by preserving the total number and lengths of all fragments in one species (*affinis*) but randomly positioning them in a genome of the same size and contig structure. I then computed the nearest neighbor distance distribution in the same way as above and fed this simulated data through the same model fit pipeline.

Best fit values for all parameters, AIC, BIC, and LRT statistics and p-values for all species and the negative control are in Table S2.

### 7. Hidden Markov Model for tinkering locus identification

To identify tinkering loci within the rest of the genome based on their lower mean distances between dispersed fragments, I modeled the genome as a two-state Hidden Markov Model (Fig. 2B) with Bernoulli emissions. Each base position *t* along a contig is assigned a hidden state S_t_ = {HOT, COLD}. These terms are used for brevity: “hot” regions correspond to tinkering loci and their higher rate of dispersed duplication, and “cold” regions correspond to the rest of the genome. Conditional on this state, a dispersed duplication of a fragment occurs with a per-base probability of *λ*_1_ (hot) or *λ*_2_ (cold), leading to a binary observation of a fragment or lack of one beginning at the position. The hidden state evolves as a first-order Markov chain with per-base transition probabilities p_(stay,hot)_ and p_(stay, cold)_. Each contig was treated as an independent run of the process.

I fit the four parameters *λ*_1_, *λ*_2_, p_(stay,hot)_, p_(stay, cold)_ to the genome-wide nearest neighbor distances of merged dispersed fragments (Methods-5, Methods-6) by maximum likelihood using the Baum-Welch algorithm ^88^. I treated the initial state distribution not as a free parameter, but as the stationary distribution of the current transition matrix at every iteration, on the rationale that the beginnings of contigs are not fundamentally different than the rest of the genome. Forward-backward and Viterbi routines were implemented in Python using just-in-time compilation via Numba^89^. *λ*_1_, *λ*_2_ were initialized as the values resulting from the best fit to the double exponential model (Methods-6), p_(stay,hot)_ as 0.9997, and p_(stay,cold)_ as 0.99998, corresponding to modest tinkering locus sizes of ∼3 kb and larger intervening regions of ∼50 kb, estimates based on observation of manually identified loci (e.g. Figure 1). Optimization stopped at convergence, defined as a change in the log-likelihood of less than 10^-6^, or at a maximum of 1000 iterations. After fitting, the hidden state of each base was inferred by posterior decoding^90^ with a threshold of 0.5. I ran the HMM on all species described in the manuscript as well as the fragment-randomized negative control (Methods-6). This gave a list of genomic regions classified as tinkering loci in each species.

### 8. Transcription quantification of genomic features

I mapped the male and female *affinis* Iso-Seq reads to the genome using minimap2^91^, with parameters -a -x splice:hq --splice-flank=no --eqx --secondary=no. I then used the genome annotations described above (Methods-4), in GFF format, to count what percent of features in each of the three categories included in the annotation (conserved genes, gene fragments, transposons) had at least one read mapped to the feature, using featureCounts ^92^ with parameters --fracOverlapFeature 0.5 --primary. Each feature (exons for conserved genes; the unmerged form of the GFF for fragments; the two genomically-based annotations of transposons (using RNA-derived annotations would be tautological)) was counted separately. Given the dataset size of a total ∼10M reads (=transcripts), about 5M per sex, this corresponds to a threshold for transcription of about 0.1 TPM in both sexes or 0.2 TPM in one sex.

This threshold corresponds to a low expression level, calling for a control to determine whether it is meaningfully above background. I used unannotated length-matched (important due to the requirement of 50% overlap of a read over the feature to produce a count in the featureCounts command) regions for this control. I created a complementary GFF containing regions not annotated as any of the above categories using BEDtools ^93^ complement, and then sampled 20,000 random regions from it, matched to the lengths of 20,000 randomly selected gene fragment and conserved gene features. I then ran the same featureCounts command as above on this unannotated region GFF. This control analysis showed that the threshold used here shows meaningfully higher expression of all categories of annotated elements above background, supporting the choice as biologically relevant. I opted for this lowest possible threshold, given that it can nonetheless distinguish noise from signal, because it maximizes sensitivity and permits capture of tissue-specific transcripts, which will necessarily be at low abundance given that RNA was extracted from whole flies. This is especially important given that new genes have been shown to be more tissue-specific than conserved genes ^11^.

### 9. Quantification of *affinis*-specific proto-genes

The goal in this analysis was to identify transcribed fragments that encode newly-formed proteins (“proto-genes”) corresponding to pieces of the conserved proteins from which they were derived. Newly formed here means created since *Drosophila affinis* diverged from its closest related species 3 million years ago. I also aimed to analyze the other fragments contained on the transcripts of these novel potential proteins. This second point is of interest for two reasons. First, it gives a sense of the degree of modular combinations of protein-coding fragments made available at tinkering loci, which speaks to the ultimate evolutionary potential of these regions. Second, protein fragments that are not coding in the context of this transcript may nonetheless have functional roles, as there is increasing evidence that protein-coding sequences also often have regulatory roles (e.g. RNA stability, RNA localization)^52^.

I performed the below analysis separately for fragments within and outside of tinkering loci.

#### 1. Identifying reads containing one or more fragments within/outside tinkering loci

I first identified Iso-Seq reads (Methods-3) that overlapped at least one fragment within/outside of a tinkering locus as determined by the HMM (Methods-7). I began with the bam of the mapped reads generated above (Methods-8); filtered out supplementary alignments with samtools view - F 2048 to remove potential PCR fusion artifacts; extracted fragments within targeted regions; and used bedtools intersect -wb -split on this GFF and the filtered bam file to generate a list of all GFF features overlapped by each selected transcript.

#### 2. Clustering reads by order and identity of fragments (“feature isoforms”)

The goal of this step was to identify what fragment(s) are present on a read. This was for the purpose of a) later determining whether they encode a protein, and b) classifying transcripts not just by their encoded proteins, but by the other protein-derived sequences that they contain (above). I used a custom script to determine, for each read, what gene fragments it contains, and in what order, 5’ to 3’. I then clustered reads that share these ordered combinations of features. These can be thought of as “feature isoforms”: reads derived from the same locus that have the same combination and order of fragments. (Note that a single “feature isoform” encompasses finer-scale variants, including start site variations, termination site variations, and alternative splicing not affecting fragment inclusion/exclusion, which deviates from the standard meaning of isoforms. The goal here is merely to more coarsely classify transcripts by their combinations of fragments.) Based on visual inspection of read alignments in IGV ^48^, there appears to have been mild 3’ degradation of some transcripts; I thus cannot exclude the possibility that this produced some purely truncated alternative feature isoforms. This is one reason that I more prominently report the numbers of loci rather than the number of feature isoforms in the main text, and aggregate feature isoforms into supersets when reporting the number of fragments per transcript.

#### 3. Identifying reads encoding fragment-derived proteins

I then used the HMMER ^85^ package’s esl-translate utility with option -m on only the sense strand of each transcript to identify all intact sense ORFs on a transcript. I used these in combination with the above clusters of “feature isoforms” via a custom script to identify reads containing an intact ORF(s) whose amino acid sequence includes sequence derived from the gene fragments on its sense strand: that is, reads transcribed from protein fragments which could produce proteins whose sequences are derived from those fragments. Where one “feature isoform” encoded multiple ORFs, either due to polycistronic transcripts or to sequence variation between the reads within the feature isoform, I then split the clusters of “feature isoforms” by the ancestral protein source(s) of their ORF(s). This formed the list of potentially protein-coding tinkering locus transcripts and the other fragments included on them.

#### 4. Identifying *affinis*-specific proto-genes

The goal of this step was to determine which of these protein-coding transcripts is specific to *Drosophila affinis* relative to all four outgroup species. The high-level approach was to search for each fragment contributing to the ORF in the genomes of all four *affinis* group outgroups (*athabasca, algonquin, azteca, helvetica)* and retain only coding fragments absent from all four outgroups, maximizing the probability of gain in *affinis* compared to the alternative of many independent losses.

The optimal method for this aim depended on the nature of the coding sequence.

#### 4a. Singleton and composite transcripts

For transcripts containing an ORF derived from only one fragment (“singleton” and “composite” in the language of the main text), the main available signal to determine the presence or absence of the coding fragment in outgroups is synteny: whether a similar sequence exists at the corresponding genomic location in the outgroup, defined by the identity of the conserved genes on either side. Because fragments are definitionally derived from ancestral genes, a standalone homology search for the fragment’s sequence is insufficient: each fragment is expected to be present in at least one copy in the ancestral gene from which it is derived. Further, many ancestral genes have dispersed several fragments that now lie throughout the genome, such that only determining whether a fragment exists at a locus other than the ancestral one, or detecting changes in copy number, is not as reliable. I therefore developed a pipeline to search for these fragments primarily by synteny.

##### 4ai. Synteny-guided protein search

First, the conserved genes on either side of the fragment (which I will refer to as “anchor” genes) in the *affinis* annotation are identified. Next, it is determined whether the two anchor genes are neighbors in the outgroup. If so, these putative syntenic regions are searched for the fragment. This is done first by analyzing the outgroup’s annotation to determine if there is a gene fragment with an annotation identical to, or in the same cluster of genes with detectable homology (Methods-4), as the annotation of the *affinis* fragment. (This is meaningfully conservative in that it excludes all *affinis*-specific duplicates of ancestral fragments within the same syntenic region, of which there are large numbers. This is likely a significant source of proto-gene undercounting. I opted for this trade-off because a) even if this stage of the pipeline retained bona fide syntenic duplicates, it would be difficult downstream to determine which are the “new” copies, making the conservation analysis unreliable, and b) analysis of additional fragments on the new protein-coding transcripts would be uncertain. Additionally, since dispersed duplicates have been shown to have a higher rate of becoming functional, and one reason for this is the difference in regulatory environments, I reasoned that these syntenic fragment duplicates were not as biologically meaningful, and so a more meaningfully accurate count could be had by excluding them here.)

Occasionally, full-length high-identity duplicates conserved in the outgroup are stochastically misannotated as the conserved ortholog. To prevent this from interfering with successfully identifying the fragment as conserved, this step also considers synteny conserved if a conserved gene in the outgroup matching the fragment’s annotation (or another in its cluster) is found on either side of one of its anchor genes. This is again a conservative choice that may reduce the true positive rate with the benefit of minimizing false positives.

##### 4aii. Synteny-guided nucleotide search

If the above synteny search fails, it could be because of a failure of annotation, due either to misidentification of the protein annotation (still strictly possible despite the cluster check above, which looks for any detectable homology between the annotations) or because the sequence has diverged too far from the *melanogaster* protein to be identified (all *affinis* group species share the same divergence time relative to *melanogaster*, but differences in substitution rate between them, and stochastic fluctuation over the detectability threshold, are possible.) In this case, the syntenic region is also searched by BLASTN, using the *affinis* fragment sequence as a query, with a very permissive E-value cutoff of 0.5. Because *affinis* is only ∼3 My diverged from its closest two outgroups, it is fairly unlikely that these nucleotide sequences will have diverged beyond detectability^19^.

##### 4aiii. Expanded synteny-guided annotation and nucleotide searches

For fragments with only one anchor gene (at the ends of contigs), the two possible syntenic regions, between the sole anchor and the genes on either side of it (reflecting the agnosticism about on which side of the anchor the fragment lies in *affinis*), in the outgroup species are searched, with both the annotation and nucleotide approaches above. This is also done for fragments with two anchor genes in *affinis* that are not found as neighbors in the outgroup, for each of the two anchors separately. This is intended to account for the possibilities that a) a local rearrangement has occurred, moving the two anchor genes apart from one another or deleting one, or b) that there is a difference in the annotation between the focal species and the outgroup that prevents the identification of the two anchors as the same.

##### 4aiv. Genome-wide nucleotide searches

It is next determined whether there is ambiguity about successful identification of the syntenic region in the outgroup species which may have prevented the fragment from being identified. For fragments not found above to be considered, if a) there are fewer than two anchor genes in *affinis*, b) the two anchor genes in *affinis* are not found to be neighbors in the outgroup, or c) the hypothesized syntenic region in the outgroup, flanked by the same two anchor genes, aligns to less than 25% of the region in *affinis* (which may suggest its misidentification), the search is expanded to look beyond synteny, searching genome-wide for an ancestral copy of the fragment. This raises the difficulties described above: there is definitionally at least one other copy of the fragment in the genome: within the ancestral gene, such that the relinquishment of synteny as the defining signal makes it difficult to distinguish which copies are orthologous. I attempted to strike a balance by searching for more local synteny, defined not by flanking conserved genes, which may be tens of kilobases away, but by the conservation of at least 3 kb flanking nucleotide sequence on at least one side of the fragment. This is again a conservative choice that risks false negatives: bona fide duplications of fragments to new genomic regions could well also copy 3 kb of additional flanking sequence. However, given the results in the main text, which finds that ectopic recombination, whose characteristic length scale is ∼2-2.5 kb, is a likely mechanism for tinkering, and that the other alternative implied by high transposon density is retrotransposition, which acts on the transcript, most of which in *Drosophila* are less than 3 kb, a 3 kb cutoff seems to strike a balance between conservatism and power. I note that this is an additional step not usually undertaken in the analysis of duplicate genes ^41^ and is likely a source of undercounting.

For this step, the entire locus in *affinis* (between the two *affinis* anchor genes, or, for genes with one anchor, on either side of the one anchor gene to the next conserved gene) is used as a query in a genome-wide minimap2 search of the target species’ genome, with parameters -g 25 -r 12 -m 0 -n 20 -p 0.1, intended to find short weak stretches of similarity. If this search reveals any region in the outgroup genome that includes both the *affinis* fragment and at least 3 kb of contiguous sequence on either side, the fragment is considered conserved in the outgroup. Minimap2 is used in preference to BLAST here because BLAST is very sensitive to small (∼50 bp) insertions and deletions, which I do not consider prohibitive for inferring a syntenic origin.

#### 4b. Chimeric transcripts

For chimeric *affinis* transcripts with multiple gene fragments that form a single intact hybrid ORF, given that each fragment exists somewhere in the outgroup genome, there is a better key signal for conservation than synteny: whether the two fragments exist anywhere in the outgroup genome in sufficient proximity and in the same relative orientation (strand and order). For these transcripts, I therefore performed a TBLASTN search using the ORF as a query with a permissive E-value threshold of 0.5, and determined whether any locus in the target genome contained all fragments (defined as covering at least 25% of the *affinis* fragment) that are a) within 10 kb of one another (a permissive distance allowing for the possibility of very long introns), b) in the same relative order, and c) on the same strand. I excluded transcripts whose different fragments fell into the same cluster of related proteins, potentially representing diverged pieces from a single ancestral gene and not true composites. (These were not homogenized in the annotation process, Methods-4, because they are not contiguous on the genome, but are on the transcript; this is not implausible because they may reflect subsequent dispersal events that moved new fragments into introns of dispersed genes, preventing contiguity but nonetheless representing the same ancestral gene rather than a chimera.) I manually excluded one chimeric protein composed of two mitochondrial proteins which are directly adjacent in *Drosophila* mitochondrial DNA, presumably missing from the outgroup assemblies.

I performed these analyses separately on each of the four outgroup species genomes, and considered a fragment *affinis*-specific if it was not detected in any.

For each transcript isoform with coding potential, I stored the length of each read’s ORF, for later use in comparing the total number of new genes to estimates from other studies (Methods-10).

Finally, I calculated the number of unique ORFs in each of the above categories; the number of unique ancestral genes that contribute fragments to each fragment isoform; and collapsed isoforms from the same genomic locus, defined as sharing at least one fragment, to count the number of unique loci.

### 10. Comparison of gene birth rates between *affinis* and other species

I use Zhou et al ^41^ to compare the rate of gene birth in *melanogaster* to that I find in *affinis* because it uses broadly similar criteria for new gene identification. It begins with annotated genes from FlyBase; the paper does not specify a release number, but the contemporaneous annotation (from 2007; https://wiki.flybase.org/wiki/FlyBase:Gene_Model_Annotation_Guidelines#Gene_Model_Annotation_Guidelines_282007.29) includes as proteins those for which there is transcriptional evidence, homology to another protein, and an ORF length of ≥50aa. I explicitly use the first two of these criteria (Methods-9), and >90% of the ORFs in my final dataset are ≥50aa (Methods-9). I report the numbers without this filter in the main text and Figures, but deduct this 10% difference in the estimates of relative rate. The paper then uses synteny and BLASTN to look for conserved genes in outgroup genomes; does not consider transcriptional evidence in outgroups; and uses five outgroups (*yakuba, simulans, sechellia, erecta, ananassae).* As described above (Methods-9), this is a similar workflow. The timescales of the comparisons are also similar. Zhou et al. analyze genes born in *Drosophila melanogaster* relative to sister species *Drosophila sechellia/simulans,* for which a large phylogenetic analysis of *Drosophilidae*, Russo et al. 2013 ^36^, estimate a divergence time of 4 million years; the same study estimates a divergence time for *affinis* and *athabasca/algonquin* of 3 million years.

There are methodological differences which are important to note. I use a more permissive E-value threshold, intended to err on the side of calling more genes as conserved (0.5 vs 10^-5^). I also implement an extra step to find the fragment elsewhere in the genome if synteny-guided search fails, which looks genome-wide and considers the fragment conserved if it finds the fragment plus 3 kb of sequence on either side intact (Methods-9). As described above, this is meaningfully conservative, potentially excluding many bona fide new duplicates. However, it also enables analysis for the nontrivial number of fragments with incomplete synteny available (at the ends of contigs, or on their own contig) due to the comparatively low assembly continuity of these genomes compared to the *melanogaster* genome; this does not strike me as obviously confounded in one direction or another, but is a clear asymmetry that could plausibly have unobvious effects. Finally, my analysis here is less conservative in using four comparator outgroups (*athabasca, algonquin, azteca, helvetica)* instead of the five in Zhou et al. Zhou et al. report a total of 72 new genes born between *melanogaster* and *sechellia/simulans,* but deem 2/72 genes to be *de novo* originated, as opposed to derived from duplication or shuffling, due to a lack of detected similarity to any other annotated genes; I remove these from their count. The majority (59/70) of the genes are tandem duplicates, which I exclude here due to the focus on hub-like tinkering loci, and therefore cannot make a meaningful comparison against. Comparing only the two rates of dispersed gene birth, Zhou et al. find a rate of 70-59=11 in 4 million years, or 2.75/million years. *affinis* tinkering loci alone have created 184 genes in 3 million years, about 20x faster than the genome-wide *melanogaster* rate; including the rest of the *affinis* genome, the total is 257 in 3 million years, a rate about 31x higher. Comparing the *affinis* rate of only new dispersed genes to the rate of all new genes in *melanogaster* removes the most common category of gene birth only in *affinis* and thus inflates the relative *melanogaster* rate significantly; it should therefore be considered a very conservative lower bound, but still yields a 5x higher rate in *affinis*.

Zhou et al. nominally report 21 chimeric genes, but use a different definition of chimeric gene than the one here. They include not only cases in which the new coding sequence includes coding sequence from parental genes, but in which it includes *noncoding* sequence from parental genes (e.g. UTRs, introns, intergenic sequence). They report 14 chimeras that “recruited … into their protein coding regions … exons.” I tried to reproduce this result by downloading the transcript sequences of all 14 genes from FlyBase, finding all intact ORFs on the sense strand, and performing a protein homology search against the *melanogaster* proteome to confirm that sequences from more than one other *melanogaster* protein were included. I was unable to reproduce the chimeric status for all but 3/14 proteins: CG12592, CG18853, and CG18217. The latter two of these do each have homology to two different *melanogaster* proteins, but in both cases these two are themselves paralogs, making a more parsimonious explanation that these are third family members; I removed these cases from my own dataset out of caution, and so do not count these to increase commensurability, resulting in 1 chimeric gene matching the definition used here. As an illustrative example of the difficulty in interpreting the meaning of chimeric genes with exons incorporated into the CDS in this study, *quetzalcoatl* is included in this list, but was the subject of a more detailed subsequent study, where it is clearly described as having a CDS that includes the sequence of one parental gene, and a UTR, not included in the CDS, with sequence from another ^94^; this matches the results of my analysis. I am unsure as to the source of this discrepancy. One possibility is that the FlyBase sequences for these transcripts have been updated since their 2008 study; another is that the intended meaning of chimera was different than interpreted here. In any case, I list the total rate of chimeric gene birth as 1 per 4 million years to match the definition used in this analysis. This is 5x lower than the rate in *affinis* (4 in 3 million years). However, the single chimeric gene in Zhou et al. is made from two tandem duplicates (CG12592, made from tandemly duplicated components of *slender lobes* and *CG18545,* which flank it directly up- and down-stream), which would therefore be excluded here. I thus do not report the 5x figure in the main text, as the rate of comparable chimeras in Zhou et al. is zero.

Because of the large uncertainty resulting from the zero count at this short evolutionary distance in Zhou et al., I also compare the rate of chimeric gene birth to that reported by Conant et al. ^43^, which sampled more events by using the much longer timescale of the divergence between *melanogaster* and *C. elegans.* They use similar methods as here, using a BLAST search of protein sequences in one species to identify proteins in the other species which cannot be found in a single contiguous hit, and are instead contained in different ancestral proteins. One difference is that they appear to rely on existing protein annotations. This may reduce the number of bona fide chimeric genes that they query, as some may have been missed; on the other hand, it may reduce the inflation of chimeric genes that they call as new if bona fide homologs are present in outgroups but unannotated ^20^. They find 48 chimeric proteins in this evolutionary interval. I could find no published studies reporting both the divergence time between *affinis* and *athabasca/algonquin* and between *melanogaster* and *elegans*, introducing the uncertainty associated with comparing divergence times across studies. TimeTree.org ^71^ reports the median estimate across many studies as 700 million years, with a range of 555-941. In the main text, I use 600 million years, on the conservative end, producing a rate of 0.08 chimeras/million years, compared to *affinis*’ rate of 1.33/million years, which is 17x faster, with a range, produced by using the above range of divergence times, of 15-26x.

## 11. Tinkering flux analyses

The first step in assessing tinkering flux per locus was to identify the loci orthologous to each *affinis* tinkering locus in outgroup species. This is detailed separately below (Methods-12).

### 1. Determination of the presence or absence of fragments at orthologous tinkering loci

For each *affinis* tinkering locus, I performed a BLASTN dc-megablast search with default parameters between the locus and the region(s) identified as orthologous in each outgroup. In contrast to minimap2, BLAST is sensitive to small (∼50 bp) insertions and deletions, and will thus detect the insertion or deletion of fragments at an orthologous tinkering locus. I used the result of this search and the positions of fragments within the *affinis* tinkering locus to determine whether each fragment was aligned to a region, and thus present, within the orthologous tinkering locus in the outgroup. I also did the converse: I considered each fragment in the outgroup tinkering locus, and used its position within the locus to determine whether it was present within the *affinis* locus. If a fragment fell into a subregion of the locus that was not covered by the orthologous locus assignment (above), I assumed that it was present. This is a conservative choice for the purpose of flux analyses. (For phylogenetic analysis, it is handled differently, per Methods-13 below.) Because these species are closely related, relying on nucleotide comparisons between them, rather than fragment annotations relative to more distant *melanogaster*, makes the divergence beyond detection of the fragments at levels high enough to affect the overall analysis relatively unlikely, especially at the extremely high default BLASTN E-value of 10, though it cannot be ruled out in individual cases.

### 2. Identification of absent fragments as new gains or duplicates

The code then distinguishes the duplication of fragments from the gain of new fragments. If two fragments in a species align to the same region in the orthologous locus, and no other region in the outgroup locus is aligned to either fragment, a duplication is inferred. If a region does not align at all, a gain is inferred.

### 3. Calculation of flux values

I then aggregated these pairwise results between *affinis* and each outgroup to determine in how many outgroups each *affinis* fragment was present or absent, as well as in how many other species each outgroup fragment was present or absent.

### 3a. Calculation of fragment gain flux

If there were multiple fragments derived from (annotated as) different regions of the same *melanogaster* gene (and thus not counted as duplicates per 2 above), I counted them as only a single gain event, on the conservative assumption that they were acquired in the same event. I then counted the number of fragments whose presence was at all variable across the phylogeny, e.g. was not present in all five species. To distinguish gain flux from duplication flux (below), I determined presence or absence only of any number of copies of the fragment at the locus: if one species had two copies and another had zero, I counted the fragment only once.

### 3b. Calculation of duplication flux

I then counted the number of duplicate copies of fragments present in any species at the locus whose presence was variable across the phylogeny.

### 4. Exclusion of tinkering loci with poor coverage in all outgroups

I. computed the percent of the *affinis* locus covered in the inferred orthologous region in each outgroup (Methods-12) and excluded loci for which this value was less than 50% in 4 out of 4 outgroups. Some missing data is tolerable in this context for two reasons. First, fragments are assumed to be present when the subregion within which they lie cannot be confidently found in an outgroup (above), such that missing data tends to deflate, rather than inflate, tinkering estimates. Second, just one species with a missing fragment (inferred only with high coverage) is sufficient to contribute the maximum total value to the total flux per tinkering locus, because each fragment counts only once if at all variable in the phylogeny, no matter its finer presence/absence pattern between species. This choice thus aims to strike a balance between conservatism and fullest use of available data. After exclusion, 243/424 tinkering loci remained. Only these are shown in Figure S4.

#### Note on interpretation

There is no phylogenetic inference in the flux calculation, which is merely raw variation. Loss events may contribute to flux, which is nonetheless a meaningful metric because it provides an intuition for the modular diversity generated by both gain and loss processes at tinkering loci (loss may enable new modular combinations by e.g. removing deleterious fragments from proto-genes). The phylogenetic analysis described below, by contrast, does attempt to specifically infer gains.

#### 12. Identification of regions orthologous to *affinis* tinkering loci

Identifying regions orthologous to tinkering loci even within this closely related taxon is challenging due to the properties that make them of interest: the rapid gain of sequence from elsewhere in the genome, internal rearrangement and duplication of sequence, and high repeat content. I sought an approach that was robust to these effects while avoiding the high cost of erroneously imputed orthologous regions, which could dramatically inflate the apparent rate of tinkering.

### 1. Identification of all similar outgroup regions

I first used minimap2 ^91^ to align each *affinis* tinkering locus to the genome of each outgroup species. Compared to other local alignment methods like BLAST, minimap2 can chain over short matching ‘anchor’ sequences that are separated by tens of kilobases or more, allowing alignments to extend across insertions and deletions as large or larger than the size of the fragments under study here. I used parameters - x asm20 -k 14 -w 5 -m 20 -n 2 to optimize this chaining permissiveness and the identification of anchors amid volatile insertions and deletions. This was intended to capture as much of the full orthologous locus as possible, irrespective of fragment-sized insertions and deletions, which were identified in a separate step.

### 2. Selection of one or more orthologous subregions

I then searched the list of outgroup regions that minimap2 returned as similar to the tinkering locus. In the simplest case, a single contiguous region in the outgroup covering the entire tinkering locus was found. However, in many cases, multiple loci, sometimes roughly contiguous with one another and sometimes not, sometimes overlapping query regions and sometimes not, were found as matching regions in the outgroup. In these cases, I collected all outgroup loci that *uniquely* covered a particular subregion of the *affinis* locus, representing unambiguous orthology, and considered them all to be parts of the orthologous tinkering locus. Often, these subregions were close to one another in the outgroup genome, separated by moderately large (>10 kb) insertions, or were adjacent and reoriented, both of which cause minimap2 to fail to join them. A minority of discontinuous regions corresponded to genuinely distant loci, on different contigs or separated by hundreds to thousands of kb, presumably brought together by genome rearrangements. Because I was interested specifically in the gain of dispersed fragments, as opposed to rearrangements bringing distant loci into contact, I did not consider these events in the fragment gain or duplication processes included in the phylogenetic analyses, although they are a potential source of new genes. (The rate of such long-distance rearrangements was not significantly higher in tinkering loci than in length-matched regions from outside of tinkering loci.)

### 3. Exclusion of subregions with ambiguous orthology

If two or more outgroup regions overlapped on the *affinis* tinkering locus by more than 400 bp, I did not assign any orthologous region to that part of the tinkering locus. This was a conservative choice: ambiguity in the true orthologous locus presents the possibility of significantly inflating tinkering event estimates in later steps, due to comparing regions that are not truly orthologous.

### 4. Calculation of coverage

Step 3 above resulted in many *affinis* tinkering loci having low “coverage,” the fraction of their sequence included in the assigned orthologous locus, in outgroups. I calculated this value for all loci in each outgroup.

#### 13. Phylogenetic tinkering analysis

This analysis begins with the pairwise presence/absence information comparing fragments between species, described above (Methods-11), and then combines them with a recent phylogeny of the species group ^70^ to assign a) dispersed fragment gain and b) fragment duplication/deletion events to individual branches in the phylogeny. The phylogeny, from Kim et al. 2024 ^70^, was generated from 1000 single-copy (BUSCO) orthologs from *Drosophilidae* using ASTRAL. The tree file with branch lengths was taken from the supplementary information for this study, available as described in that manuscript at https://zenodo.org/records/11200892 (Fig1_S1_data.treefile).

### 1. Branch assignment of fragment gain events

For fragment gain, I assigned gain events to the branch directly preceding the divergence of all species in which it is present. For example, if a fragment is present in both *helvetica* (the earliest-branching species) and *affinis*, even if absent in all other species, I attributed it to the branch before the root of the lineage. Fragments attributed to the branch before the root of the group were not included in any subsequent analyses, which tests only branches within the phylogeny. This approach corresponds to an assumption that convergent gain of the same fragment at the same locus is prohibitively unlikely compared to multiple losses, followed by a secondary parsimony assumption for losses (attributing a gain to any branch earlier than this requires an additional loss).

### 2. Branch assignment of fragment duplication events

For fragment duplication, I used the total count of each duplicated fragment in each species as input to the Sankoff dynamic programming algorithm for parsimony-based phylogenetic inference ^95^ to infer the number of events on each branch. This weights duplications and deletions equally, which differs from 1) above. This seems appropriate, because gains and losses for dispersed fragments are not symmetrical processes – *dispersed* fragment gain is unlikely compared to *local* fragment losses – while local duplications and deletions are much more symmetrical. I count both local duplications and deletions resulting from the algorithm as events on branches.

### 3. Handling of low-coverage loci and ambiguous data

To handle the issue of many tinkering loci not having full orthologous regions confidently identified in outgroup species, I applied an additional analysis step for fragments whose loci had coverage lower than 95%. For fragments marked present in low-coverage regions (the default when data did not allow a direct assessment of their presence or absence; Methods-11), I determined whether their presence or absence would affect the final branch assignment (which would not be the case, for example, if it were not the earliest-diverging species with present fragment in the fragment gain analysis). Only in this case, I excluded the fragment from analysis. This is a source of substantial loss of power (below), but seems to deal fairly with missing data. (As expected, I found in earlier analyses that unilaterally assigning fragments at poorly covered loci to an older or younger branch has a very large effect on the inferred rates, such that it did not seem appropriate to make such a unilateral default assignment.)

#### 4. Statistical tests for rate heterogeneity across the phylogeny

The combination of generally low numbers of events per locus, the even smaller number of events per branch, and the exclusion of fragments that could not be assigned unambiguously to a branch, per 3 above, resulted in total numbers of events per branch that were sufficiently small that there was usually not sufficient power for the most straightforward tests of statistically significant changes in rate across branches for single loci. This motivated two alternative tests.

##### 4a. Per-branch test, aggregated over all loci

I took the inferred number of a) gain or b) duplication/deletion events per branch for each tinkering locus, and summed these over all loci to find the total number of events per branch. I then used the branch lengths from the phylogeny (above) to calculate significant deviations from the total number of expected events per branch, calculated according to a null model in which fragment gains and duplications happen at a uniform rate across the phylogeny, such that events per branch are proportional to its length. This was modeled as a binomial distribution, with N = total number of observed events and p_branch_ = L_branch_ / Σ _branches_ L_branch_ (each branch’s length divided by the sum of all branch lengths, i.e. its proportion of the total length within the phylogeny). I then FDR corrected the p-value for the 8 (one per branch) tests using the Benjamini-Hochberg method ^96^, at an adjusted p-value of 0.05. I also assessed whether each branch had strictly sufficient power by calculating the hypothetical p-value in the maximally anomalous scenario in which all events were assigned to that branch. (All branches had sufficient power in this aggregated test.)

##### 4b. Global enrichment per-locus test

A second test overcomes the issue of low power in testing individual loci by using a global enrichment strategy: testing for an *excess* of significant results above a calculated expected background of false positives ^97^. The idea here is that, although no one locus may have statistical power sufficient to survive FDR correction, the total set of p-values may contain an excess of significant values compared to the expected number under the null, such that signal is apparent. Though beneficial for power, it has the downside of sacrificing the ability to identify *which* particular loci are the false positives vs true positives. This test uses the same binomial framework as above for each locus and each branch individually, using N = total number of observed events per locus and a probability per branch of that branch’s fraction of the total branch lengths in the phylogeny. For each locus, I calculated whether each branch had power to produce a p-value of less than 0.05 in the best-case (maximally anomalous) scenario where all fragments fall on that branch. A locus was considered powered overall if it had at least one such branch. I then determined whether any powered branch in the locus had a significant (two-tailed, at p=0.05) p-value by this binomial test, and considered a locus significant if it had at least one significant branch. I then calculated the number of loci expected to be significant by this definition by chance. This is not 5%, because the test is “at least one significant branch at p=0.05” over the multiple powered branches for a locus. The null probability of a specific distribution of fragments along branches is given by a multinomial with 8 components, one for each branch, with the branch-specific probability the same as in the binomial. The probability of at least one branch having a significant result under the null is calculated in the case of fragment gain by explicitly enumerating all possible combinations of branches and then summing those which give a significant p-value for one or more branches. In the case of fragment duplication, where the overall numbers of events are higher, explicit enumeration is not computationally tractable, and so it is approximated by a Monte Carlo simulation using 100,000 draws. This null probability is summed over all loci to produce the expected number of significant loci, which is compared to the observed number. The statistical significance of the observed number relative to this expectation is given by a Poisson binomial, which is approximated here by a Poisson distribution, a good proxy when the probabilities are small.

### Caveats

Because tinkering loci are structurally volatile, they likely produce many loss events. Additionally, hybridization is known to occur within the *affinis* group ^98^. These factors introduce uncertainty into phylogenetic analyses. Indeed, the phylogenetic pattern of about 30% of fragments was not strictly concordant with the species phylogeny, requiring more than one event (e.g. gain and then subsequent loss, or hybridization after gain) to be explained. The methods here assume that hybridization and fragment loss are not so common as to significantly affect results.

#### 14. Relationships between the three modes of tinkering

### Correlation between fragment flux and duplication flux

After excluding tinkering loci with coverage too low for a reliable estimate of flux (Methods-11), I calculated the Pearson correlation between the fragment flux and duplication flux for each of the remaining loci. I then calculated the p-value by a permutation test.

### Comparisons of flux between rearranged and non-rearranged tinkering loci

After excluding tinkering loci with coverage too low for a reliable estimate of flux (Methods-11), I used the mannwhitneyu function in the SciPy ^87^ stats package to perform a two-tailed Mann-Whitney U test comparing the distributions of each type of flux values (fragment and duplication) for the tinkering loci with and without rearrangements anywhere in the phylogeny. P-values are reported in the main text. The common language effect size (the proportion of the time that a value randomly sampled from one distribution would be larger than a value randomly sampled from the other distribution) is 0.7 for fragment flux and 0.83 for duplication flux.

#### 15. Rate heterogeneity calculation for locus in Figure 4

There are multiple loci in the *Drosophila algonquin* assembly corresponding to a subregion of this locus, likely due to high heterozygosity that the assembler failed to collapse. (This might be expected at structurally volatile tinkering loci, but is notable because the assembly is derived from a single animal.) For this reason, my automated pipeline did not identify an orthologous locus in *algonquin*, resulting in all fragments at the locus being called as ambiguous in their phylogenetic placement, and the rate heterogeneity at the locus thus not being calculated due to lack of power. Manual analysis revealed that all loci are orthologous and confirmed that all *affinis* fragments are absent (for example, contig JAWNKY010002279.1:1-14,426), allowing manual calculation of the p-value with the same methodology described above. The locus is a good example of the loss of data to the conservatism of the approach given the difficulties of sequence analysis at volatile tinkering loci.

#### 16. Ectopic recombination sequence analyses

### 1. Quantifying fragments that share flanking sequence with parental exons

To determine how many fragments show signatures consistent with ectopic recombination, I extracted the 2500 nt up- and downstream from the fragment and the 2500 nt up- and down-stream of the matching (sharing at least one amino acid position corresponding to the *melanogaster* protein per the annotation, Methods-4) exon in the parental gene, and performed a BLASTN dc-megablast search with parameters -dust no -evalue 0.01. I turned off dust to allow inclusion of repetitive sequences, common ectopic recombination substrates. I then determined whether there was a resulting alignment of at least 100 nt between the flanking sequences. As a control, I also performed the same search between the flanking regions of each fragment and one randomly selected exon of a conserved gene. A length of 2500 nt for up- and down-stream flanking sequences was selected to match the typical lengths of recombination tracts ^99^. A shared ≥100 nt region within this range at a permissive E-value allows degradation of the original recombination tract due to mutations like small indels and rearrangements, which may be especially likely at tinkering loci.

### 2. Identifying fragments with flanking sequences shared with parental transcripts

The alternative possibility for a dispersal mechanism is retrotransposition. In this case, the flanking region of the fragment should share sequence flanking the parental exon as included in transcripts of the parental gene. The relative explanatory power of these alternatives can be determined by assessing the rate at which the same up/downstream 2500nt of a gene fragment matches sequence at the genomic parental locus but not transcripts from the parental locus, and vice versa.

#### 2a. Identifying isoforms from parental genes

I first used the mapped *affinis* reads (Methods-8) to identify reads mapped to conserved genes according to my GFF-formatted annotation with BEDtools ^93^ intersect. I then clustered all reads mapped to each conserved gene, with MMseqs2 ^86^ easy-cluster, with parameters --min-seq-id 0.95 -c 0.3 --cov-mode 1 --cluster-mode 2, to create a set of representative transcripts, roughly corresponding to isoforms, for each gene. I also identified genes for which no transcripts were found (a minority, per the earlier assessment of the rate of transcription of genes annotated as conserved).

#### 2b. Search for shared sequence between parental gene isoforms and fragment flanking sequence

I then used samtools ^100^ to combine the GFF and the bam file for the mapped reads to extract the coordinates of each protein exon on each of its representative reads. I then performed the same BLAST search as for the genomic loci, now searching for sequence shared between the 2500 nt up- and downstream flanking sequence of each fragment and all representative transcripts from its parental gene. I excluded the parental exon itself, analogously to the genomic search (and to avoid tautological hits), by removing the read-relative coordinates of the matching parental exon on each transcript from analysis. I then determined whether there was a region of similarity at least 100 nt in length somewhere in the transcript outside of the shared exon itself.

### 3. Comparison of shared sequence tracts between parental gene transcripts and the parental gene locus

I then determined how many gene fragments discriminated between the two hypotheses, and in which direction. For each fragment, I determined a) if it shared sequence with the parental genomic locus; b) if it shared sequence with at least one parental transcript; and c) if transcripts were identified for the parental gene (meaning that a meaningful answer to b) is possible). Fragments for which a) was true, b) was false, and c) was true were counted as favoring the hypothesis of ectopic recombination. Those for which a) was false, b) was true, and c) was true were counted as favoring the hypothesis of retrotransposition. 64 sequences favored retrotransposition, and 419 favored ectopic recombination. As expected, a majority of fragments did not discriminate between the two hypotheses, either because they share sequence with both (41%) or neither (44%) the genomic locus and the transcript. The former category occurs because much flanking sequence around genomically encoded exons is also contained in transcripts, and vice versa. The latter category is because, especially for older fragments (all *affinis* tinkering locus fragments were included here, regardless of inferred age), subsequent mutations, especially the structural mutations common at tinkering loci, have likely eroded the signature of the original dispersal mechanism.

#### 17. Transposon enrichment

### 1. Overall enrichment around genomic features

The hypothesis of transposon-mediated ectopic recombination (ER) predicts that regions involved in the ER process are enriched for transposons. These regions are those adjacent to a) the gene fragments derived from ER and b) the parental exons from which they are derived. A natural set of control regions to which to compare the transposon densities in a) is c) intergenic regions that do not overlap with parental exons, non-parental exons, or any gene fragments. Because these regions are expected to have minimal function, they serve as an estimate of the “background” rate of transposon density. A natural set of control regions to which to compare b) is d) all non-parental exons: those which do not match any tinkering locus fragment. Because all conserved exons likely have some function, it is not appropriate to compare them to unannotated regions; comparing parental to non-parental exons preserves closer functional constraints, as well as a closer genomic and regulatory environment (e.g. chromatin accessibility), factors which may influence transposon density.

To select control regions (c), I randomly selected regions matched to the lengths of tinkering locus fragments within the genome; discarded any that overlapped with any conserved gene or gene fragment; and then extracted the 2500 nt up- and down-stream of it, repeating the process until a total number of control regions matching the total number of gene fragments was sampled.

To assess the transposon density around these four classes of regions, for each feature, I calculated the percentage of the 2500 nt upstream and downstream (for a total of 5000 nt) that was included in one or more features in the RNA-based transposon annotation (Methods-4.7). This resulted in a distribution of transposon density for each of the four categories of elements. I compared these distributions with a two-sided Mann-Whitney U test. These are reported in the main text.

I also performed this analysis using genomic DFAM transposon annotations (Methods-4.4a). Parental exons had significantly higher transposon density than non-parental exons (mean 2x, 0.008 vs 0.004, both medians 0, Mann-Whitney U p=2*10^-25^), and tinkering locus fragments had marginally significantly higher density than control regions (mean 1.5x, both medians 0, Mann-Whitney U p=0.07). Because the transposon density with the RNA-based annotations is much closer to existing estimates of transposon density (20% overall vs 22% here in control regions, 2-8% in easily-assembled euchromatic regions vs 3-8% around conserved exons ^101^) than with the genomic DFAM annotations (7% in control regions, 0.4-0.8% around conserved exons), I take the former result as more reliable. I interpret the lower numbers from the DFAM annotations as a result of the DFAM sequences being sourced from highly diverged (∼30 Mya) *melanogaster* transposons, which have diverged sufficiently that many are not detected, which may be exacerbated at volatile tinkering loci.

### 2. Enrichment of classes of transposons at tinkering locus fragments

To assess whether this overall enrichment was driven by particular classes of transposons, I first compared the density of different transposon types around tinkering locus fragments to those around their inter-feature control regions, and those around parental exons to those around non-parental exons, taking these two pairs to be good comparators for the same reasons as in 1) above. For each transposon “family,” represented by a clustered Iso-Seq read annotated as a transposon (Methods-4.7) with significant sequence similarity to one or more places in the genome, I performed two statistical tests. The first is a Mann-Whitney U test comparing the transposon *density* distribution in the 2500 nt flanking regions around tinkering locus fragments to the distribution in the flanking regions around control regions. The second is a Fisher’s exact test comparing the *number* of flanking regions in each category containing *at least one sequence of* the transposon (regardless of the length of the annotated region). I then FDR-corrected the resulting p-values for each test and considered transposons enriched if they had a resulting adjusted p-value of <0.05 by *both* tests.

I converted from transposon sequence families, represented by clustered Iso-Seq reads annotated as transposons, to broad classes of transposons on the basis of which transposon PFAM domains were detected in the sequence, according to the table in (Methods-4.4b). This was a coarse assignment, not intending to classify families beyond the very broadest levels, where domain presence is straightforwardly diagnostic.

#### 18. Evolutionary analysis of *Rcc1-Txl* chimera

### 1. Inference of gene origin and duplication events

I determined the evolutionary history of the chimeric gene by searching Dipteran and other genomes broadly for homologs. I searched Dipteran genomes using a combination of manual TBLASTN searches, BLASTN searches, and searches of my annotations of *affinis* group species and close outgroup *Drosophila pseudoobscura;* targeted TBLASTN/BLASTN searches specifically of the orthologous loci in *affinis* group species and closely related species *Drosophila pseudoobscura;* and TBLASTN searches against *Drosophila pseudoobscura, persimilis, melanogaster,* and all other species included in the NCBI NR (via BLASTP) and NT (via TBLASTN) databases. All BLAST searches used permissive E-value thresholds of 0.5. As expected, both individual gene fragments, from ancestral genes *Rcc1* and *Txl*, are present in *Drosophilidae* genomes, in the context of the full-length genes at their inferred ancestral syntenic locations. My annotations for *pseudoobscura* and the *affinis* group also showed additional *Rcc1* fragments (4+) dispersed through the genome in all species. However, fragments from these two genes are never within ∼150kb of one another in any species more divergent from *affinis* than *Drosophila azteca*, where they are immediately adjacent (<200 nt), at the locus orthologous (judged based on synteny to flanking conserved genes) to one of the copies of the chimera in *affinis*. This configuration, at this locus, is also present in *athabasca* and *algonquin*. This is simply explainable by the chimera’s origin in the common ancestor of *affinis* and *azteca* ∼4 Mya.

I inferred duplications as described in the main text with the same approach, using parsimony as a criterion: copies of the chimera at the same locus (judged by identity of syntenic genes) in two species were inferred to have originated from a single event in the common ancestor of that species. The resulting inferences had little ambiguity, requiring only one duplication event to be explained, except for one case: a copy that is fully intact only in *athabasca* and *algonquin*, but for which the orthologous locus in both *affinis* and *azteca* contains only one of the two constituent fragments (*Rcc1*). This is the copy shown as the first locus under Step 2 in Fig 5. This is consistent with a scenario in which a) this copy was gained on the same branch, numbered 2 in Fig. 5, as the other oldest copy, and subsequently degraded in *affinis* and *azteca*, or in which b) this copy resulted from a more recent duplication (<3 Mya) in the common ancestor of *algonquin* and *athabasca*, with convergent gains of the *Rcc1* fragment at the *affinis* and *azteca* loci independently. However, loss seems likelier than 2 convergent gains of the *Rcc1* fragment at the same locus, such that I tentatively favor a). I am unable to resolve the order in which the two copies of the chimera gained on branch 2 appeared (Fig. 5 depicts this ambiguity in that the arrows do not point to a particular locus).

### 2. dN/dS analysis

I performed the dN/dS analysis using the ancestral copy that has not degraded (second locus under Step 2 in Figure 5), as this copy had the maximum number of orthologs available, maximizing power. To perform the dN/dS analysis, I extracted the coding sequence of the *affinis* gene, the only chimeric ORF on the transcript. I then inferred the orthologous ORF in the other species. In *affinis*, the ORF is not interrupted by introns, straightforwardly encoded in genome sequence; I thus searched for intact chimeric ORFs at the genomic locus in *athabasca, azteca, and algonquin*. In the former two species, there was also an intact chimeric ORF at this locus, which I used as the input to the dN/dS analysis. In *algonquin*, although both fragments remain present and adjacent at the locus, there is no genomically-encoded intact ORF containing both, indicating either that splicing is required for an in-frame ORF or that the chimeric ORF is not conserved. I therefore used only the *affinis, athabasca, and azteca* sequences in the dN/dS analysis. I translated all three ORFs, aligned the amino acid sequences with MAFFT version 7.271 ^102^, and removed columns with gaps, resulting in 134 aligned residues. These corresponded to two distinct subregions of the proteins; the three species vary in presence and absence of blocks of indels between these regions, corresponding to different retained/deleted segments of the ancestral *Rcc1* and *Txl* genes. These variable regions were not included in the analysis and may or may not be conserved, but the remaining residues in these conserved subregions were sufficient in number for a dN/dS estimate of the selective pressure operating on them. I substituted the codon sequence into the alignment, and ran codeml m0 from PAML ^103^ to estimate a single *ω* across all three branches, using two codon frequency models, F3×4 and F61. F3×4 had a log likelihood of −781.6, a maximum likelihood *ω* of 0.48, and a 95% CI of (0.2, 0.76). F61 gave similar results and had a log likelihood of −737.6, a maximum likelihood *ω* of 0.43 and a 95% CI of (0.18, 0.69). To rule out a strong anomaly on a single branch driving this result, I also used yn00 to estimate all pairwise *ω* values, which ranged from 0.43 to 0.60.

## Data availability

Raw Iso-Seq data in FASTQ format are available from NCBI SRA under BioProject PRJNA1476202.

All other data supporting results and figures, too large to be included as supplemental information here, as well as all custom code used here, are available at https://github.com/caraweisman/Weisman_tinkering_loci_2026.

## Supplemental Information

### Supplemental Figures

**Figure S1:**
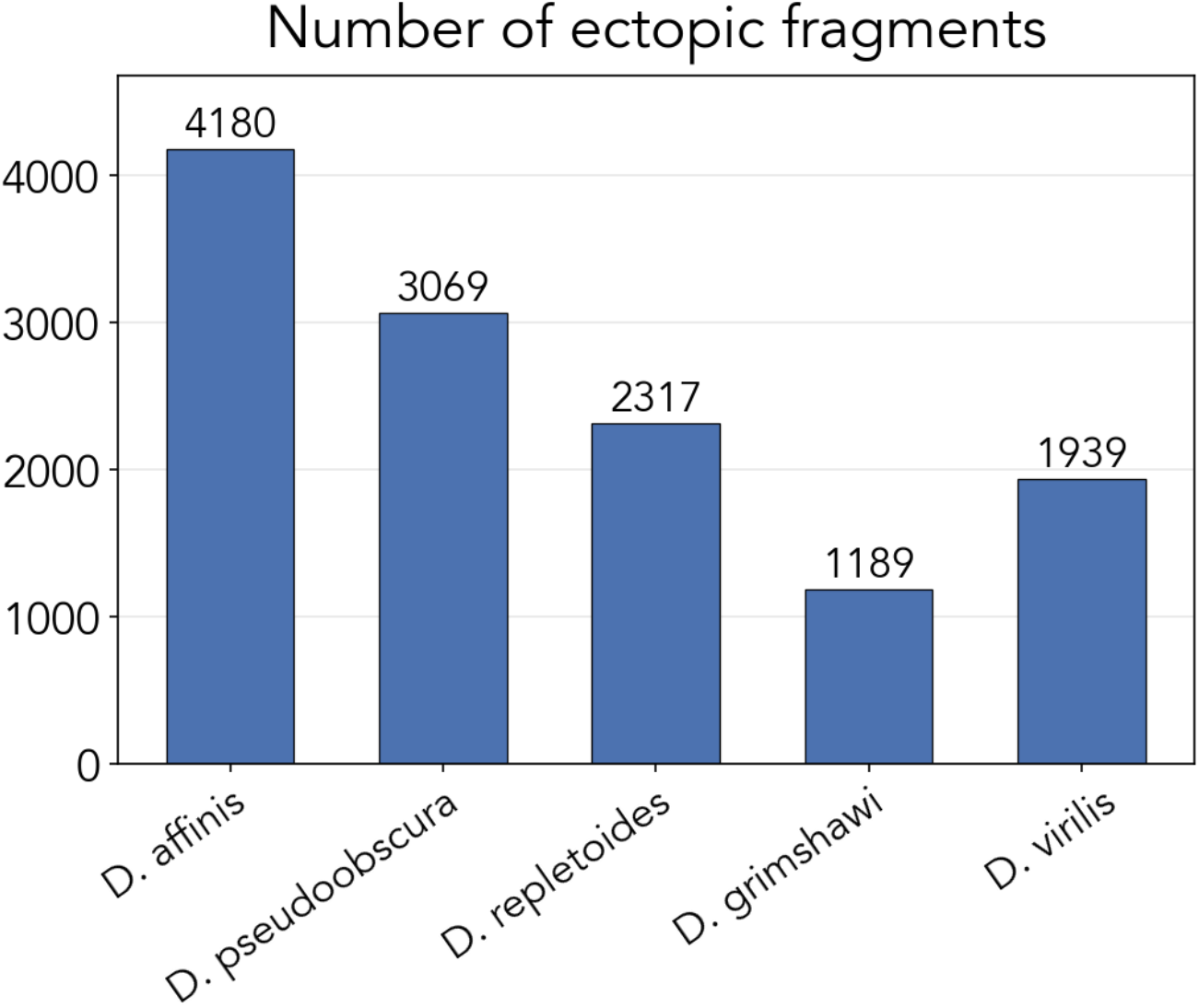
Number of dispersed fragments in each of the five *Drosophila* species.

**Figure S2:**
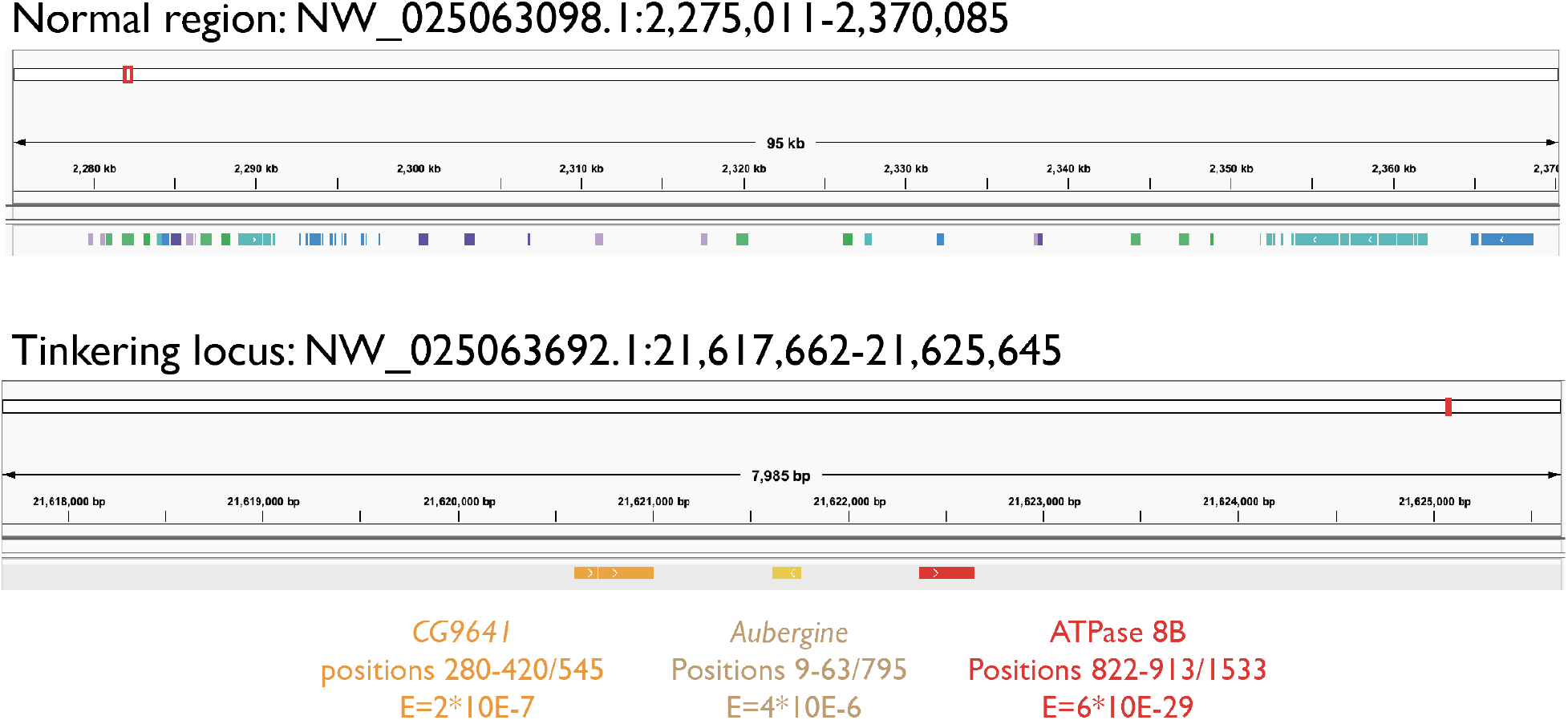
Examples of a region not identified as a tinkering locus (top) and a region identified as a tinkering locus (bottom) by the HMM from *Drosophila grimshawi,* as visualized in IGV ^48^ using the GFF-format annotations produced here. Fragments are annotated in warm colors (red/orange/yellow) and conserved genes in cool colors (green/blue/purple). Names, positions relative to the *melanogaster* homolog, and E-values relative to the *melanogaster* ortholog of features are provided for the tinkering locus but not for the normal region due to the size of the latter, but are available in the GFF and/or via IGV.

**Figure S3:**
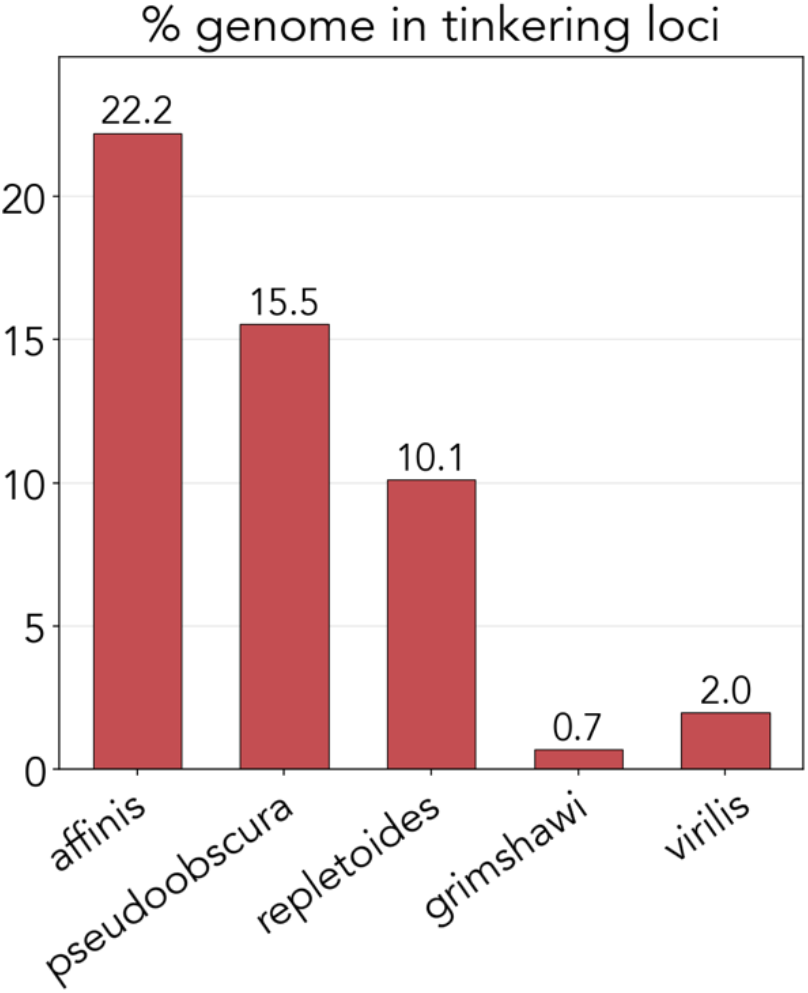
Total percent of genome contained in tinkering loci for each of the five phylogenetically broad *Drosophila* species.

**Figure S4:**
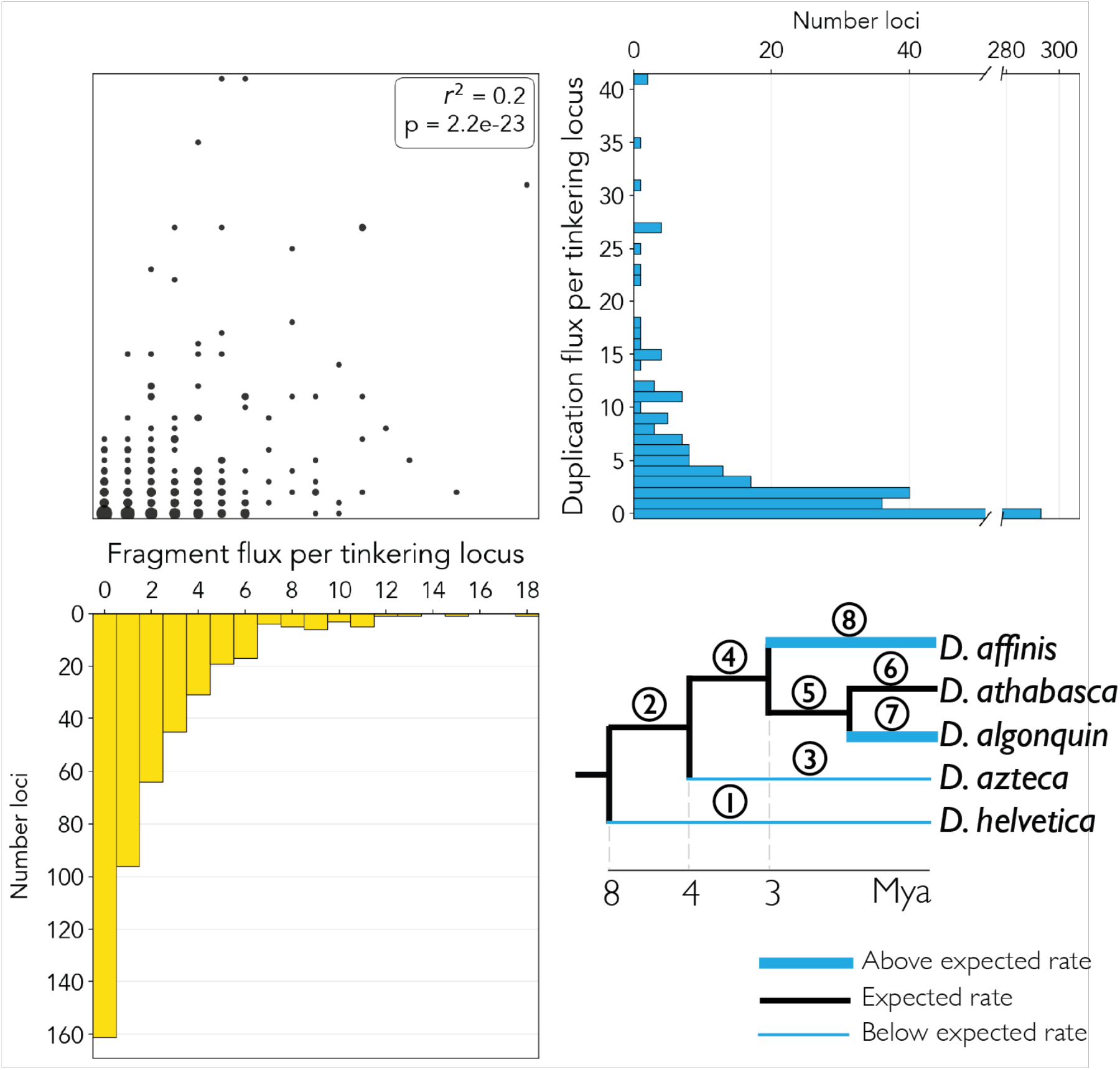
Duplicate flux and phylogenetic duplication analysis. Top left: correlation between the duplication flux (y axis) and fragment flux (x axis) of individual tinkering loci. Top right: marginal histogram of duplication flux over tinkering loci. Bottom left: marginal histogram of fragment flux over tinkering loci. Bottom right: phylogenetic analysis of changes in duplication rate over the *affinis* group phylogeny, aggregated over all tinkering loci, as in Fig. 3C.

**Figure S5:**
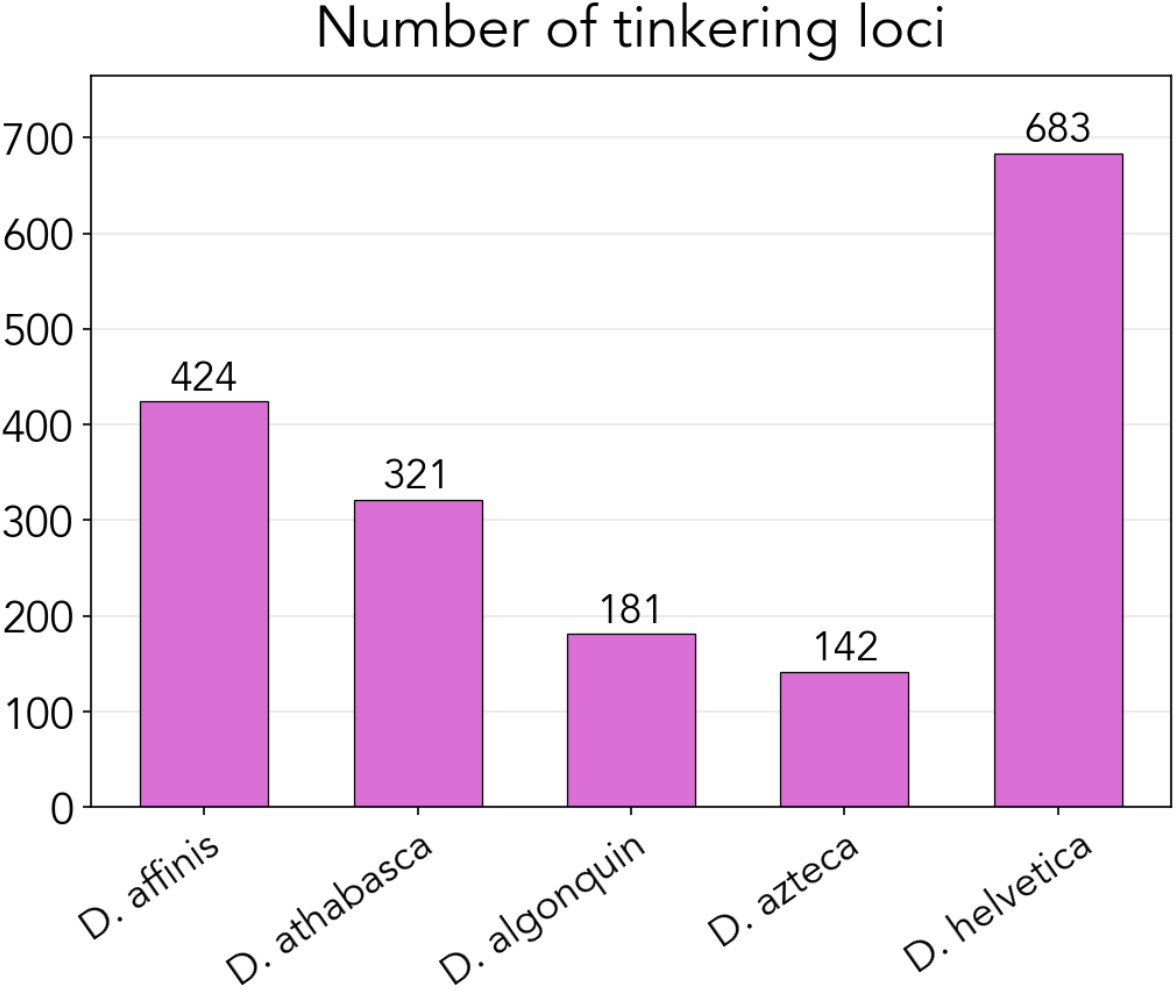
Number of tinkering loci in *affinis* group species.

### Supplemental Tables

**Table S1:** RefSeq/Genbank accessions of all genome assemblies used here.

| Species | NCBI<br>(Refseq/Genbank)<br>accession | Sequencing<br>platform(s) | Long-read? | Notes |
| --- | --- | --- | --- | --- |
| *Drosophila affinis* | GCA_035045985.1 | Oxford<br>Nanopore;<br>Illumina | Yes |  |
| *Drosophila athabasca* | GCA_008121215.1 | PacBio | Yes |  |
| *Drosophila algonquin* | GCA_035041765.1 | Oxford<br>Nanopore;<br>Illumina | Yes |  |
| *Drosophila azteca* | GCA_005876895.1 | PacBio Sequel;<br>Illumina | Yes |  |
| *Drosophila helvetica* | GCA_963969585.1<br>(primary assembly),<br>GCA_963969575.1<br>(secondary assembly;<br>used for second<br>helvetica chromosome<br>in Figure 1) | PacBio, Arima2 | Yes | Phased assembly |
| *Drosophila pseudoobscura* | GCF_009870125.1 | PacBio Sequel | Yes |  |
| *Drosophila repletoidea* | GCA_018150835.1 | Oxford<br>Nanopore<br>Minlon; Illumina<br>HiSeq | Yes |  |
| *Drosophila grimshawi* | GCF_018153295.1 | Oxford<br>Nanopore<br>Minlon; Illumina<br>HiSeq | Yes |  |
| *Drosophila virilis* | GCF_030788295.1 | PacBio RSII | Yes |  |
| *Chymomyza caudatula* | GCA_035041775.1 | Oxford<br>Nanopore;<br>Illumina | Yes |  |
| *Chymomyza procnemis* | GCA_035046065.1 | Oxford<br>Nanopore;<br>Illumina | Yes |  |
| *Scaptodrosophila inornata* | GCA_054122035.1 | Illumina Hi-Seq | No |  |
| *Drosophila sucinea* | GCA_018150745.1 | Oxford<br>Nanopore<br>Minlon; Illumina<br>HiSeq | Yes |  |
| *Drosophila paulistorum* | GCA_018152135.1 | Oxford<br>Nanopore<br>Minlon; Illumina<br>HiSeq | Yes |  |

**Table S2:**
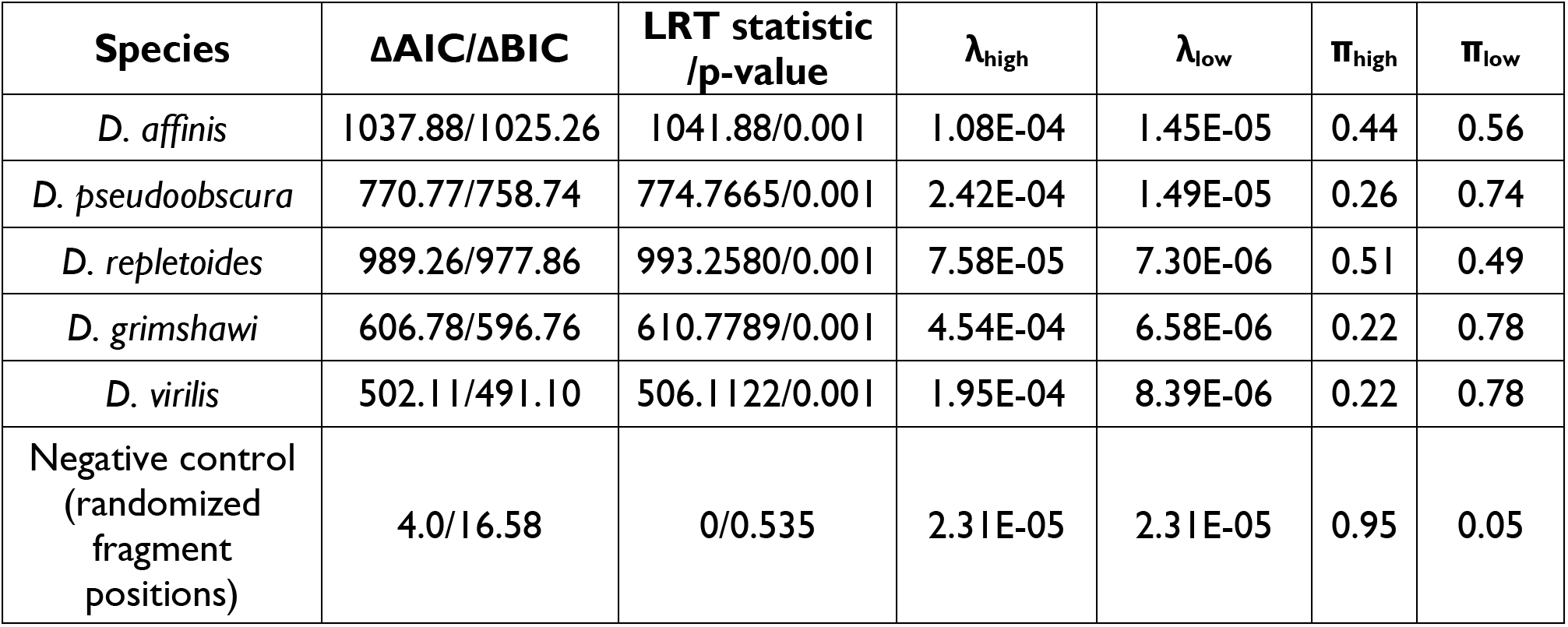
For all species plus the negative control: LRT p-values and statistics, AIC, and BIC values for fits to single and double exponentials; best fit lambda and pi values for the double exponential.

| Species | $\Delta\text{AIC}/\Delta\text{BIC}$ | LRT statistic<br>/p-value | $\lambda_{\text{high}}$ | $\lambda_{\text{low}}$ | $\pi_{\text{high}}$ | $\pi_{\text{low}}$ |
| --- | --- | --- | --- | --- | --- | --- |
| *D. affinis* | 1037.88/1025.26 | 1041.88/0.001 | 1.08E-04 | 1.45E-05 | 0.44 | 0.56 |
| *D. pseudoobscura* | 770.77/758.74 | 774.7665/0.001 | 2.42E-04 | 1.49E-05 | 0.26 | 0.74 |
| *D. repletoidea* | 989.26/977.86 | 993.2580/0.001 | 7.58E-05 | 7.30E-06 | 0.51 | 0.49 |
| *D. grimshawi* | 606.78/596.76 | 610.7789/0.001 | 4.54E-04 | 6.58E-06 | 0.22 | 0.78 |
| *D. virilis* | 502.11/491.10 | 506.1122/0.001 | 1.95E-04 | 8.39E-06 | 0.22 | 0.78 |
| Negative control<br>(randomized<br>fragment<br>positions) | 4.0/16.58 | 0/0.535 | 2.31E-05 | 2.31E-05 | 0.95 | 0.05 |

**Table S3:** Transcription statistics for the three annotation feature categories (conserved genes, fragments, transposons) and unannotated negative controls.

| Annotation category | % transcribed<br>($\geq 1$ read) | Mean<br>(TPM) | 5%<br>(TPM) | 25%<br>(TPM) | 50%<br>(TPM) | 75%<br>(TPM) | 95%<br>(TPM) |
| --- | --- | --- | --- | --- | --- | --- | --- |
| Conserved | 96% | 63 | 0.1 | 2.1 | 10.8 | 41.8 | 215.5 |
| Tinkering locus fragments | 56% | 14 | 0 | 0 | 0.1 | 1.5 | 35.4 |
| Transposons | 25% | 0.15 | 0 | 0 | 0 | 0.1 | 0.57 |
| Intergenic controls<br>(length-matched) | 8% | 0.04 | 0 | 0 | 0 | 0 | 0.01 |

**Table S4**: List of singleton ORFs, chimeric ORFs, and composite ORFs, with read counts, TPM, contigs, and position, for tinkering loci and outside tinkering loci. (Attached as separate file.)

**Table S5**: Results of flux analyses and rate heterogeneity tests for fragment gain and duplication. (Attached as separate file.)

**Table S6**: Results from statistical tests for transposon family enrichment. (Attached as separate file.)

## Notes

### Competing Interest Statement

The authors have declared no competing interest.

### Summary of Updates

Abstract and discussion altered for stronger emphasis on key takeaways thanks to feedback; clarifications and elaborations of points throughout the results.

